# The origin and evolution of cultivated rice and genomic signatures of heterosis for yield traits in super-hybrid rice

**DOI:** 10.1101/2024.03.19.585738

**Authors:** Daliang Liu, Hao Yin, Tao Li, Liang Wang, Song Lu, Houlin Yu, Xinhao Sun, Taikui Zhang, Quanzhi Zhao, Yiyong Zhao

## Abstract

Unraveling evolutionary history and genomic basis of heterosis is fundamental for advancing rice productivity. We developed a genome-scale phylogeny of *Oryzeae* by coalescing 39,984 gene trees. Our analysis supports parallel, independent origins and nearly synchronous evolutionary trajectories leading to the subsequent domestication of *indica* and *japonica*, evidenced by molecular dating and synonymous substitution rates for syntenic and domestication-associated genes. Our survey of 1,383 gene duplications in ancestor of *O. sativa* uncovers their roles in vital biological processes, highlighting the significance in environmental adaptability. Additionally, we confirm the lack of hybridization event among subspecies ancestors through gene tree topology and D-statistical analyses. We generated 71.67 GB whole-genome sequencing data for five super-hybrid rice varieties and their progenitors, revealing differential positive selection and genetic exchanges between subspecies, essential for heterosis formation. Crucially, our study underscores the role of non-additive gene expression in heterosis, particularly in genes associated with DNA repair and recombination, which may confer resistance traits. Furthermore, eQTL and de novo mutation analyses identify key developmental and stress response genes, offering targets for enhancing heterosis in rice. Overall, our research reveals crucial insights into the genetics of rice domestication and heterosis, offering a genomic resource to improve rice’s agricultural productivity.

## Introduction

Rice (*Oryza*), along with maize and wheat, is one of the three principal global cereal crops and is cultivated as a primary crop in diverse environmental settings. Rice originated approximately 130 million years ago in Gondwanaland, the disintegration of Gondwanaland led to speciation across different continental plates with varied genome types (AA, BB, CC, BBCC, CCDD, EE, FF, GG, HHJJ, and HHKK), including 20 wild species and two cultivated varieties, namely Asian rice (*Oryza sativa* L.) and African rice (*Oryza glaberrima*) ^1,2^. Despite the generally poor agronomic traits of wild species, varieties from different growth regions exhibit distinct stress resistance and serve as reservoirs for excellent natural genetic resources ^3–5^.

The high yield of Asian cultivated rice has led to its widespread cultivation globally, providing approximately 20% of the human caloric intake and serving as a primary food source ^6^. It is especially popular in Asian countries such as China, India, Japan, and Korea, where over 90% of rice cultivation and consumption occurs, providing the main source of carbohydrates and energy to billions of people ^7,8^. Previous studies have suggested that Asian cultivated rice can be primarily classified into *indica*, *japonica* (tropical *japonica* and temperate *japonica*), *aus*, and aromatic subgroups ^6^. Genetic, ecogeographical, and archaeological perspectives have revealed substantial differences between *indica* and *japonica*, which represents the two main subspecies of Asian cultivated rice ^2,9^. The differentiation between temperate and tropical *japonica* originated from adaptive evolution in distinct regions, with *aus* being more closely related to *indica*, whereas aromatic characteristics are closely associated with *japonica* ^10^.

Despite many research on Asian cultivated rice, because of its economic importance and status as a model crop organism ^2,11–15^ and modern domesticated rice is a product of repeated cross-breeding, and the issue of the origin of the subgroups of cultivated rice has yet to be resolved. Two prevailing hypotheses exist regarding the origins of Asian cultivated rice: a single origin of *indica* and *japonica* and multiple origins. The single origin hypothesis suggests that the two main cultivated subspecies, *indica* and *japonica*, were domesticated from the wild species *O. rufipogon*. For example, studies of the *prog1* gene, which controls upright growth, and the *Bh4* gene, which controls hull color, suggest a common origin for *indica* and *japonica* ^16,17^.

Subsequent population analyses based on single nucleotide polymorphisms (SNPs) from 630 genes support the single-origin domestication of rice ^18^, as well as the hypothesis that *japonica* originating from the wild rice *O. rufipogon* in southern China formed *indica* by migrating and merging with local wild resources in south and southeast Asia ^19^. In contrast, multiple origin hypothesis suggests that Asian cultivated rice was independently domesticated from wild rice of different lineages. For example, Genome-wide comparative analyses of the nuclear genomes from wild rice species *Oryza rufipogon* accessions W1943 and W0106, along with those of *indica* cultivars, in conjunction with studies of chloroplast genomes derived from diverse rice populations, have led to the postulation that *indica* and *japonica* subspecies underwent independent domestication events ^20,21^. Corroboratively, another study also proposed independent domestication of *indica*, *japonica*, and *aus* ^10^. More recent research indicates that rice domestication likely began independently from different wild strains in various regions, followed by the continuous selection and exchange of beneficial alleles among different cultivated varietal groups ^22^. The origin of cultivated rice remains a topic of considerable debate. In general, the evidence supporting a single-origin hypothesis predominantly stems from studies examining individual or small number of genes. The domestication and origins of Asian cultivated rice are exceptionally complex, and the reliance on single or partial genes, as well as singular research methodologies, would fails to provide a comprehensive and reliable evidence to indicate the evolutionary history of cultivated rice.

Heterosis is a phenomenon in which the offspring (F1) resulting from crosses between genetically different parents exhibit phenotypic traits superior to those of the parents. This phenomenon is widely applied in agricultural production and is considered a hallmark of innovation in modern agriculture. Heterosis plays a pivotal role in improving the yield and quality of various crops with significant economic implications ^23,24^. The molecular genetic enhancement of heterosis, particularly within the context of rice cultivation, has been significantly advanced following the initiation of the “Chinese Super-Hybrid Rice Breeding Program.” This strategic program has facilitated the development of numerous superior hybrid rice varieties, leveraging the genetic diversity and heterosis phenomenon to improve yield, resistance to pests and diseases, and overall crop resilience. The representative of elite hybrid rice varieties include Liangyoupeijiu (LYP9), Y Liangyou 1 (Y1), Y Liangyou 2 (Y2), Y Liangyou 900 (Y900), and Xiang Liangyou 900, also known as Chaoyou 1000 (CY1000). At the turn of the 21^st^ century, LYP9, derived from the combination of male parent Yangdao 6 (R9311) and female parent Peiai 64SA (PA64SA), achieved a yield potential of 9.75∼10.5 t/hm2 ^25^; whereas Y1, a second-generation super-hybrid rice, retained the superior male parent Yangdao 6 and introduced the photoperiod-thermo-sensitive male sterile line Y58S with *japonica* lineage as the female parent, enhancing the utilization of intersubspecific heterosis and resistance, with a yield potential reaching 12 t/hm2 ^26,27^. The cultivation success of Y2 marked its entry into the third-generation of super rice ^28^, with improvements in the sink (organs for volume and nutrient absorption), source (organs for the production and export of photosynthates), and flow (internal transport system), achieving a yield potential of 13.9 t/hm2 ^29^. The development of Y900 represents a milestone achievement in the pursuit of the fourth-generation super rice objective. Distinct from its progenitor strains, Y900 was derived utilizing an intermediate type of restorer line, R900, which combines the genetic backgrounds of both *indica* and *japonica* subspecies, as the male progenitor, while Y58S was selected as the female progenitor. This breeding strategy effectively harnesses the intersubspecific heterosis of *indica* and *japonica* rice varieties. Phenotypically, Y900 exhibits advantageous agronomic traits, including augmented foliar thickness, an elongated period of photosynthetic activity, and enhanced resistance to lodging. These attributes collectively contribute to superior yield potential of Y900, which is estimated to reach up to 15.0 t/hm2 ^30^. Subsequently, the salt-tolerant variety CY1000, characterized by lodging resistance and high yield, achieved an impressive peak yield of 18.0 t/hm2 ^31^. In summary, the genetic mechanisms behind the generational improvement of hybrid rice yields remain elusive. Successive successes in breeding high-yielding super rice varieties have provided excellent material for elucidating the genomic signatures underlying the formation of yield trait advantages.

Heterosis is underpinned by three main classical hypotheses: dominance (partial and complete) ^32^, overdominance ^33^, and epistasis ^34,35^. In addition, gene expression patterns analysis on heterozygous hybrid F_1_ in maize also supported the above three classical heterosis hypotheses ^36^. Subsequent research has demonstrated the occurrence of partial dominance, complete dominance, and overdominance effects across a multitude of genes, as evidenced by comprehensive transcriptomic analyses. For instance, the *SFT* gene in tomato ^35^, *Dw1*, *Dw3*, and *qHT7.1* genes in sorghum ^37^, and *GS3* and *Ghd7* genes in rice ^38^ have been demonstrated to contribute to phenotypic superiority through dominance or overdominance effects. Recent advancements in genomics, when combined with high-throughput sequencing technologies, have facilitated a deeper understanding of the molecular mechanisms that underpin heterosis. Transcriptomic analyses have led to a deeper understanding of the relationship between allele-specific expression and hybrid dominance. Studies have shown that for specific loci in hybrid rice, either allele of the parent may be more advantages under different spatiotemporal conditions, allowing hybrids to selectively express the beneficial allele ^39^. The ’homo-insufficiency under an insufficient background’ model suggests that homozygote functionality insufficiency under insufficient backgrounds (environment and regulatory factors) is central to the non-additive effects (dominance and epistasis) and the formation of heterosis, theoretically demonstrating that non-additivity and the formation of hybrid dominance are inextricably linked to genetic backgrounds ^40^. Collectively, these comprehensive datasets, derived from the aforementioned studies, present an invaluable opportunity for dissecting and understanding the underlying genetic mechanisms responsible for the expression of dominance traits in these cultivars.

Phenotypic polymorphisms are frequently correlated with underlying genetic variation. Genes act as critical molecular intermediaries that facilitate the transduction of genetic diversity into observable phenotypic traits, thereby contributing significantly to phenotypic heterogeneity within species. Gene expression modulation is primarily governed by *cis*-regulatory elements situated in close proximity to the gene of interest ^41^. Additionally, trans-acting factors, such as transcription factors, and various metabolites, which may exert their regulatory influence from a remote locus, play a pivotal role in the dynamic regulation of gene expression ^41^. These regulatory mechanisms collectively orchestrate the intricate expression patterns that are essential for the development, adaptation, and evolution of organisms ^41^. The analysis of expression quantitative trait loci (eQTL), which correlates quantitative traits with variant loci via linear regression, effectively establishes a link between phenotypic polymorphisms and genetic variation, which is crucial for dissecting the genetic architecture of complex traits ^41–43^. The development of next-generation sequencing technologies has led to an increase in eQTL studies. For instance, a study on human populations has identified regulatory loci for phenotypes such as hair color, shape, and thickness ^44^, and analysis of more than 30,000 blood samples has revealed potential drivers for more than 1000 phenotypes ^45^. Studies of the early development of maize kernels have identified grain-size-related genes ^46^. These eQTL related studies enhanced our understandings of the regulatory mechanisms in animals and plants, providing significant insights into how genetic variations affect gene expression and species phenotypes.

In this study, we aim to reveal the origin of cultivated rice and the genomic signature of heterosis for yield traits in super-hybrid rice by integrating multi-omics data. Our study executed a comprehensive genome-scale phylogenomic analysis, coupled with divergence time estimation, and assessed evolutionary rates of pivotal domestication genes alongside gene duplication events. This was accomplished by incorporating 33 high-quality genomes from Oryzeae, which comprised two genetically distinct lineages of Asian wild rice, as well as closely related genera *Leersia* and *Zizania*. The integration of these genomes enabled a robust comparative framework, facilitating a deeper understanding of the genetic and evolutionary underpinnings that have shaped the domestication processes within this complex group of grasses. In addition, we conducted comprehensive analyses to elucidates the hybridization and gene flow dynamics among the most recent common ancestors (MRCA) of *indica* and *japonica* rice varieties, as well as among elite super-hybrid rice varieties. These analyses encompassed the detection of hybridization events through the application of phylogenetic invariants, which emerge under the coalescent model incorporating hybridization. Furthermore, we employed phylogenetic network approaches to identify migration events and conducted population structure analysis to further understand the genetic relationships and flow patterns among rice varieties. Collectively, our study provides evidence for the retention of ancient genetic lineages from diverse rice varieties in cultivated rice, indicating possible hybridization in ancestral *Oryza* species. Contrarily, our data do not support extensive hybridization between the progenitors of the *indica* and *japonica* subspecies. Instead, it appears that the *indica* and *japonica* subspecies may have arisen from two independent domestication events involving different wild rice populations.

In this study, we executed whole-genome sequencing on five leading *indica* hybrid rices (LYP9, Y1, Y2, Y900, and CY1000) and their progenitors, covering a wide range of genomic variations. Transcriptomic data from three super-hybrids (CY1000, Y900, and LYP9) and their parental lines were integrated with mutation profiles for eQTL analysis, aiming to elucidate the genetic complexities associated with their superior agronomic traits. The mutation analysis indicates a prevalence of non-synonymous mutations in super-hybrids, denoting strong positive selection in these super-hybrids. Apart from Y1 and CY1000, the other super-hybrids shared more SNPs with their paternal lineage, indicating a paternal genetic influence. Hybridization detection and population genetics analyses provided strong evidence of extensive hybridization and introgression between the five commercialized super-hybrids and *japonica* rice, hinting at the incorporation of varied genetic resources in the cultivation of super-hybrids rice. A divergent gene expression pattern among hybrids highlighted the potential importance of non-additive genes in trait dominance. Collectively, the analyses of eQTL and de novo SNPs identification revealed numerous key genes related to yield, offering insights into the genetic architecture that drives the yield advantage in super-hybrid rice.

In summary, this research provides a robust phylogenomic framework that enhances our understanding of rice domestication, underpinned by a comprehensive analysis of gene flow and hybridization in rice species. Our novel insights into the genetic basis of yield superiority in super-hybrid rice varieties underscore the intricate evolutionary dynamics that have been harnessed to drive agricultural innovation and productivity.

## Results

### Large-scale phylogenomic profiles support the independent origins of *indica* and *japonica* subspecies

From archaeological and genetic evidence various contradictory scenarios for the origin of different varieties of cultivated rice have been proposed ^10^. The origin and domestication of the Asian cultivated rice *indica* and *japonica* have been the focal points of research, with some prevailing controversies. Herein, we reconstructed the phylogenetic relationships among *Oryza* spp. At the genomic level, selecting 33 high-quality genomes across various rice subgroups in Oryzeae (Table S1) along with the outgroup *Brachypodium distachyon* genome. Our study generated an extensive set of 39,984 gene family trees, which coalesced into a highly supported species tree. Notably, members of both *indica* and *japonica* were distinctly separated into two monophyletic groups (Fig. 1a), underscoring the independent origins of *indica* and *japonica*. Our data further revealed that *O. rufipongon* w1654 and *O. rufipongon* w1943 represent the most divergent lineages within the *indica* and *japonica* subgroups, respectively. These findings are in alignment with previous studies that propose perennial wild rice *O. rufipongon* and annual wild rice *O. nivara* as the domestication ancestors of *O. sativa* ^2,19^. Intriguingly, our data also revealed that annual *O. nivara* is the sister group to the *aus* subgroup. Furthermore, the combination of *O. nivara* and *aus* emerges as the sister to the *indica* group, consistent with prior research ^10^. This comprehensive phylogenomic profiling thus supports the independent origins of the *indica* and *japonica* subspecies.

**Fig. 1.**
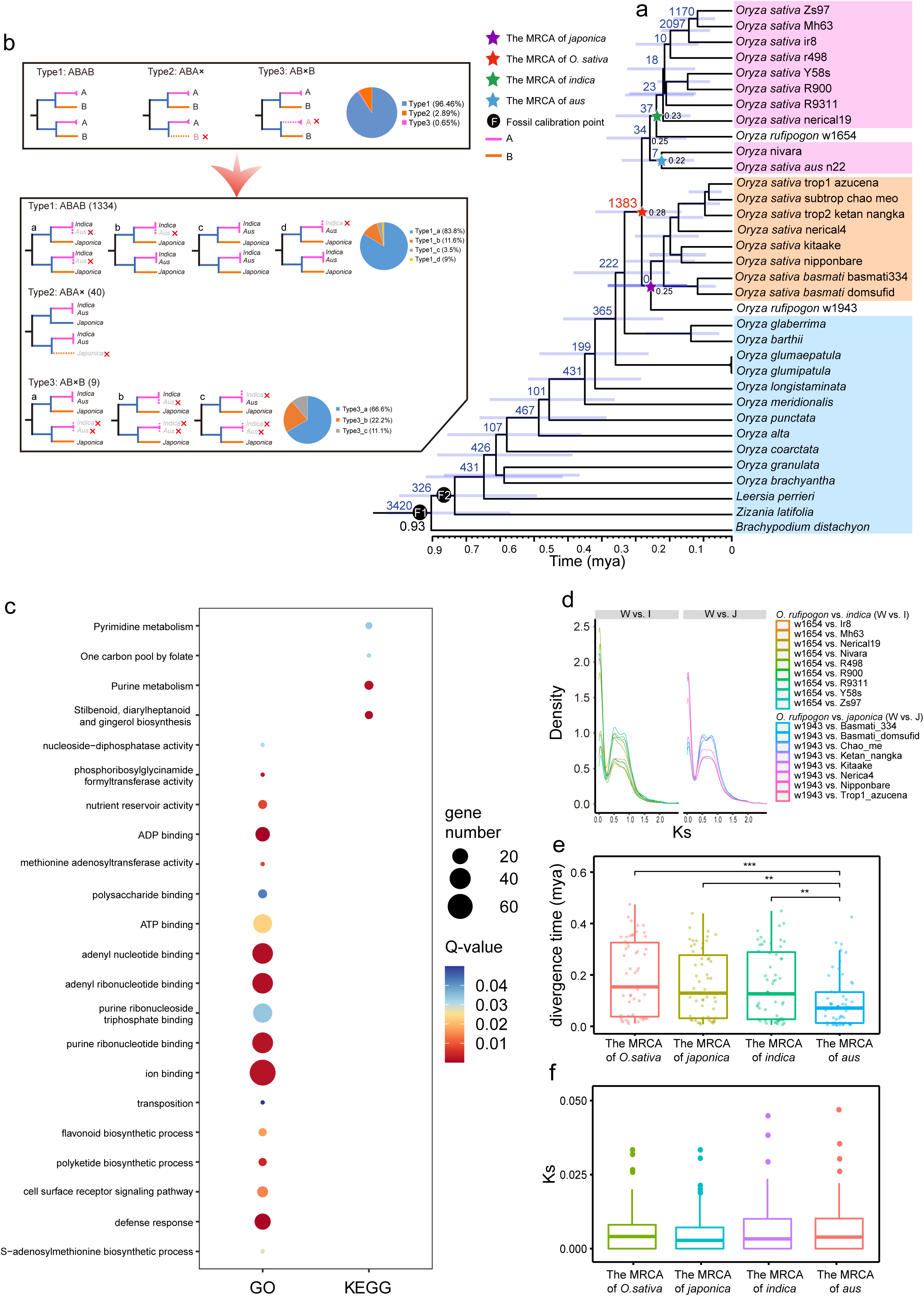
Genome-scale phylogenomic analysis and divergence among *Oryza* species. (a) Phylogenetic tree depicting the phylogenetic relationships among various rice varieties in the context of geological times. The blue bars on the node of species tree represent the 95% highest posterior density (HPD) for the estimated divergence times that all nodes in the species tree have support values greater than 0.95. The pentagrams represent the MRCA of each rice subgroup, with divergence times denoted to the right. The number of gene duplication events is indicated at the upper left of each node, with the red star marking the MRCA of *Oryza sativa*. (b) Topological classification of the duplication event in the MRCA of the *Oryza sativa* into three types, illustrated by different patterns (ABAB, ABAX, and ABXB) with corresponding pie charts showing the proportion of each type, to elucidate the evolutionary history among Asian rice subspecies. (c). This figure aims to elucidate the significance of the genes associated with the duplications of the MRCA of *Oryza sativa*, from the perspectives of biological processes (BP) in the Gene Ontology (GO) and key pathways in the Kyoto Encyclopedia of Genes and Genomes (KEGG) (Q-value < 0.05), highlighting their importance for Asian rice species. (d) Intraspecific all syntenic gene pairs Ks density profiles of wild rice versus cultivated rice species.(e) Boxplot displaying the divergence time estimates for 54 putative domestication genes from the regions of low nucleotide diversity in the MRCA of *O. sativa*, *japonica*, *indica*, and *aus* rice subgroups. (f) Boxplot showing the distribution of Ks values for 54 domestication genes across the same ancestral nodes of Asian rice subgroups as in (e).

The presence of Ks peaks from paralogous genes within a genome could serve as an indicator of gene duplication burst events (Fig. S1b). To further analyze these events, we utilized Tree2gd ^47^ to conduct a phylogenomic profiling of gene duplication events across all gene family trees. This analysis led to the identification of a total of 1,383 gene duplication events at the most recent common ancestor node of *O. sativa*, as depicted in Fig. 1a. The gene tree topologies for these 1,383 gene duplications were categorized into ABAB, ABA, and ABB types (Fig. 1b), accounting for 96.46%, 2.89%, and 0.65% of the total, respectively. The dominant prevalence of gene duplications retained in the ABAB type suggested the absence of a hybridization signal among the MRCAs of the *indica*, *japonica*, and *aus* subspecies groups.

To further clarify the mechanism underlying the 1,383 gene duplication (GD) events observed at the ancestral node of Asian cultivated rice (Fig. 1a), we conducted comprehensive syntenic dot plot and Ks analyses for Nipponbare (representing the *japonica* subgroup) and R9311 (representing the *indica* subgroup). The absence of a syntenic block with a smaller Ks value negates the possibility of a recent whole genome duplication event in the MRCA of *Oryza sativa* (Fig. S2c-d). In the genomes of the representative species Nipponbare and R9311, the retained gene duplication events from the aforementioned 1,383 GDs involved 402 and 81 gene duplicated pairs, respectively, which exhibit weak collinearity (Table S6). Specifically, Nipponbare and R9311 account for 0.25% and 1.23% of the gene duplication events with syntenic evidence, respectively (Fig. S2a-b). Furthermore, the proportions of tandem gene duplication events in Nipponbare and R9311 account for 3.23% and 20.98%, respectively (Fig. S2e-f, Table S6). In summary, tandem duplication significantly contributes to the occurrence of gene duplication events in the MRCA of *Oryza sativa*. Additionally, a total of 359 and 218 unique genes under gene duplication events were identified in the MRCA of *Oryza sativa* in Nipponbare and R9311, respectively (Table S7). A Chi-square test for genes with transposable elements (TEs) within a 2 kb region upstream and downstream of the coordinates revealed a significant correlation (P-value < 0.05) between the duplicated genes and TE distribution. This suggests that TE insertions may also contribute to the origin of the gene duplication burst at the ancestral node of cultivated Asian rice. Collectively, these findings support the notion that these GDs did not arise as a result of allopolyploidy. This is consistent with the pattern that the majority of GDs in the cultivated rice ancestor were ABAB topology type, suggesting that the cultivated rice ancestor likely did not undergo a hybridization event amongst different subspecies ancestors. However, it is plausible that there may have been a minimal number of gene introgression or gene flow events.

### Large-scaled gene duplicates retained from the MRCA of *Oryza sativa* may contribute to environmental adaptation

Gene duplication events are critical in providing raw material for the evolution of gene novelty and complexity. In the MRCA of *Oryza sativa*, a comprehensive reconciliation of gene family trees with the species tree revealed a total of 1,383 gene duplication events (refer to Fig. 1a). Following identification, the 158 genes derived from the above 1,383 duplication events and retained within the *Oryza sativa* Nipponbare genome were subjected to functional annotation and pathway enrichment analyses. The analyses employed GO and KEGG databases to elucidate the functional implications of gene duplications derived from the MRCA of *Oryza sativa*. Our results revealed a total of 18 enriched GO terms, comprising 12 molecular functions (MF), and six biological processes (BP) terms (Fig. 1c, Table S2), indicative of the diverse roles these gene duplicates may play. Notably, biological processes such as the defense response (GO:0006952) and cell surface receptor signaling pathway (GO:0007166) were significantly enriched. The defense response encompasses the array of physiological and biochemical reactions by an organism to counteract potential threats, including pathogens, herbivores, or environmental stressors, thereby safeguarding their health and integrity ^48,49^. The cell surface receptor signaling pathway represents a crucial mechanism for plant responses to external stimuli, both biological and abiotic ^50^. Furthermore, pathway enrichment analysis highlighted the importance of three specific pathways: the stilbenoid, diarylheptanoid, and gingerol biosynthesis pathway (ko00945), along with Purine metabolism (ko00230). Stilbenoids have been recognized for their role in conferring resistance to rice blast disease, whereas diarylheptanoids are implicated in plant defense against herbivory^51^. These findings underscore the critical function of gene duplicates from the MRCA of *Oryza sativa* in defense mechanisms, which could be instrumental in bolstering the resilience of rice populations and their adaptive capacity to environmental challenges.

### The molecular clock and Ks analysis of orthologous and domesticated genes reveal parallel and nearly synchronous evolutionary trajectories leading to the domestication of *indica* and *japonica*

Understanding the domestication history of rice is crucial for the conservation and utilization of germplasm resources for advanced breeding. We estimated the divergence time of rice species using molecular clock analysis (Fig. 1a). The molecular clock results indicated that the divergence time of the most recent common ancestor of each subspecies of cultivated rice followed the order: *Oryza rufipogon* w1943 > *Oryza rufipogon* w1654 > *Oryza nivara*. During speciation, gene sequences typically accumulate random nucleotide variants at a certain base mutation frequency. An analysis of the Ks density distribution of orthologous gene pairs between species in each subpopulation and their recent common ancestor revealed that Ks in the *indica* and *japonica* combinations exhibited similar distribution trends and peaks (Ks values ranging from 0.5 to 1) (Fig. 1d). For a more comprehensive comparison, we displayed boxplots of the Ks values of the MRCA of the *japonica* and *indica* subspecies in combination with the subspecies representative varieties Nipponbare vs *O. rufipogon* w1943 and R9311 vs *O. rufipogon* w1654, respectively. (Fig. S1a). This clearly showed that the Ks values of the *indica* and *japonica* groups were similar, while the *aus* group was the smallest. This suggests that the *indica* and *japonica* subspecies diverged and formed at similar times from their ancestral species formation, while the *aus* may formed much later. Compared to natural selection, domestication expedites the stabilization of the specific trait for certain populations, thereby reducing nucleotide diversity in the corresponding sequence regions ^52^. Among the previously reported nucleotide low-diversity regions located in the *japonica*, *indica*, and *aus* subgroups ^10^, we screened 54 putative domestication-associated genes across all species (see Methods). To assess the relative timing of domestication and to mitigate potential systematic errors introduced by concatenating multiple gene sequences, we conducted molecular clock analyses on the 54 OGs. This was done by concatenating all genes as well as analyzing each gene individually. Subsequently, we performed boxplot statistics on the domestication times of individual genes. The results indicated that there was no significant difference in the domestication time between *indica* and *japonica* rice, and the *aus* values were relatively small (Fig. 1e, Fig. S1c, Table S4). Meanwhile, the results of the Ks distribution of 54 putative domestication-associated genes in different subgroups showed that the Ks values were similar in the *indica* and *japonica* groups, but the *aus* group did not show a consistent trend out of the molecular clock (Fig. 1f, Table S5), In general, the onset of domestication was similar in the MRCA of *indica* and *japonica* subspecies.

The existence of gene flow between *indica* and *japonica* subspecies implies that they are not simply completely independent modes of domestication ^42^. We conducted and analysis of Ka and Ks for 11 well-known rice domestication genes, as reported in previous studies ^42,53–55^ between species within the *indica*, *japonica* subgroups and their recent common ancestral subspecies to characterize the domestication origin of these genes in *indica* and *japonica* subspecies (Fig. S3). The results indicated that fluctuations in the Ka/Ks values of some genes suggest that the domestication process has been subjected to varying degrees of positive selection. Some genes exhibited similar Ka and Ks values within subgroups but significant differences between them, such as *GW5* ^56^, *PROG1* ^57^, and *OsC1* ^58^, which were associated with grain shape, plant morphology, and pericarp luster. The Ks values of these genes were significantly higher in the *japonica* subpopulation than in the entire *indica* subpopulation, indicating that they originated and domesticated independently in the *indica* and *japonica* subpopulations, with *japonica* preceding *indica*. Some genes showed consistent Ka and Ks values in the *indica* and *japonica* subgroups, suggesting that they may have begun domestication in both subspecies simultaneously or as a result of gene flow during the early divergence of the subspecies, such as *SDR4* ^59^, *SH4* ^60^, *SHAT1* ^61^ and *WX* ^62^, which were associated with seed dormancy, seed shattering, and glutinous endosperm. In summary, these results lead us to hypothesize that subspecies formation occurred through different stages of evolution and that for some of the key traits, *japonica* began to be domesticated earlier than *indica*. In the subsequent history, in both subspecies, some traits were noticed and domesticated concurrently.

### Analyses of hybridization support ancient introgression from early divergent species of Oryzeae may contribute to rice speciation, with limited evidence support a hybridization origin for each MRCAs of *indica, japonica* and *aus* subspecies

To further investigate the possibility of hybridization events between the ancestral nodes of *indica* and *japonica*, we utilized HyDe ^63^ to analyze hybridization and introgression signals. For HyDe analysis, we both considered orthologous genes and genes derived from gene duplication at the MRCA of *Oryza sativa*. We established two gene sets for hybridization analysis. The first set is a supermatrix derived from the concatenation of 669 orthologous groups (OGs) with a coverage rate of 50% for *Oryza sativa* samples in the species tree of this study and all these single-copy orthologous genes were split and pruned from the 1383 gene duplication (GD) events that retained in the MRCA of *Oryza sativa*. The second dataset comprises a supermatrix generated by concatenating 1,300 orthologous groups (OGs) with 100% taxa coverage, representing orthologues across species within our phylogeny (Fig. 1a and Fig. S4). The two datasets previously mentioned were employed in hybridization analysis to clarify the ancestral state of *Oryza sativa* and assess the extent of genetic introgression. Our study of orthologous genes (OGs) retained from duplicated genes at the MRCA of rice revealed no hybridization signals for the MRCA of *indica*, *japonica*, and *aus* subpopulations (Fig. S4 a, c, e and g). However, introgression analysis involving 1,300 orthologs demonstrated introgression signals from the *japonica* subpopulation to *indica* (Fig. S4b, d, f, and h), indicating that *japonica* genetics were incorporated into *indica* subgroups post the divergence of their common ancestors (Fig. S4d). This finding aligns with the practices of modern hybrid breeding, where genes from japonica are introduced into indica through hybridization, illustrating a historical precedent for current breeding strategies. In addition, in our study, the ancestors of the two major subgroups of cultivated rice contained genetic signals from ancient wild rice of the genus *Oryza* (Fig. S4 b, c, d and f), suggesting that that introgression from ancient divergent species of Oryzeae may have played an important role in facilitating the speciation of cultivated rice. Meanwhile, these wild rice species including BB (*Oryza punctata*), and CCDD (*Oryza alta*) types of wild rice genomes, a result which broadens previous suggestions that genetic introgression between AA types of wild rice exist in African wild rice. genetic introgression between only *Oryza glumaepatula* and *Oryza barthii* in African wild rice ^64^.

Furthermore, to explore the relationship between *indica* and *japonica* rice within Asian cultivated rice, we examined the introgression relationship between the *indica* and *japonica* subspecies in the analysis of 1,300 single-copy OGs. As in previous studies ^20,22^, We found a large amount of contributing genetic material between the *japonica* and *indica* subpopulations (Fig. 2a,b) suggesting that genetic introgression between these two subspecies is very frequent through model hybrid rice breeding procedures. On the other hand, we analyzed some *japonica*, *indica*, and wild rice genomic next-generation sequencing (NGS) data collected from previous studies (Table S8) to obtain 35,502 high-quality SNPs and conducted a genetic network analysis of these species’s SNPs using Treemix ^19,65^, and simulations were run with 0-8 migration events (each migration event was repeated five times). Based on the previous described method, the best number of migrations (m) was estimated among populations in the dataset ^66^, with results show that the variance explained and likelihood values stabilize at Δm = 3 (Fig. 2c), suggesting that three migrations are the best fit. Specifically, observations included migration signals from the ancestral *indica* to *japonica*, gene flow from *japonica* to *aus*, and migration from *indica* to *japonica*, indicating shared genetic material among subtypes.

**Fig. 2.**
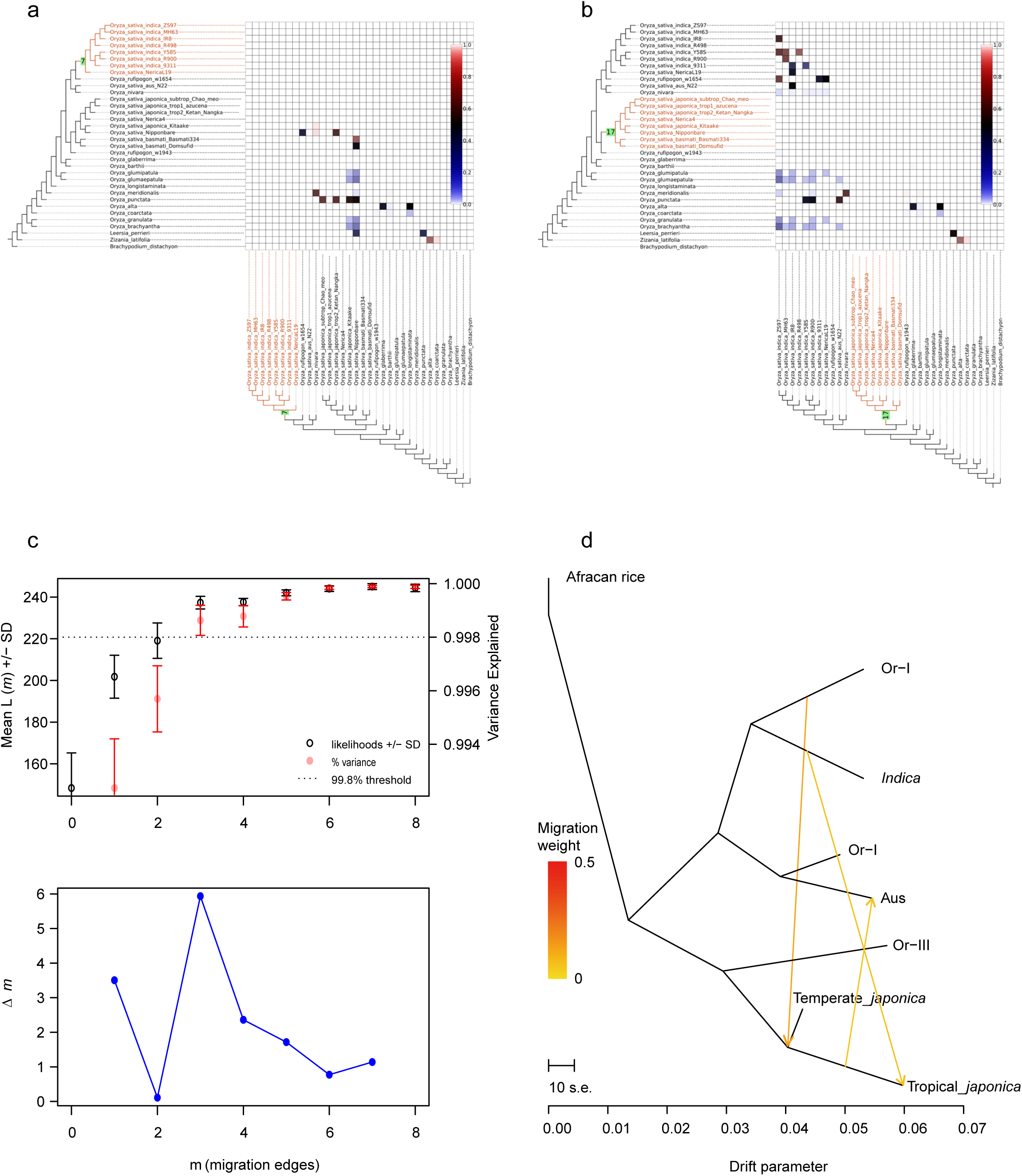
Hybridization, introgression, and migration dynamics across Asian cultivated rice subpopulations. (a-b) The heatmap illustrates the hybridization signals detected by the HyDe software based on 1,300 single-copy gene retained at the MRCA of the indica, japonica and aus subspecies, annotated with red-highlighted branches on the cladogram to denote the detected clade. Each small colored square on the heatmap corresponds to a hybridization signal identified by HyDe, with the color intensity representing the probability of inheritance for the associated taxa on the y-axis. Contiguous squares of identical shading that compose a larger block within the heatmap, along with their corresponding taxa on the axes forming a monophyletic clade, suggest the MRCA of that clade as a potential parental lineage for the MRCA of the studied clade, which is indicated with a node number on a green background. (c) The graph illustrates the optimal migration influx ascertained using the OptM package, with Δm indicating the identification of three significant migration events. (d) A phylogenetic network inferred by TreeMix, outlines the pattern of migrations between subpopulations. The arrows, annotated in colors proportional to the ancestry contributions, direct from the contribution (donor) to receiving (recipient) subpopulation, thus mapping the route of gene flow.

Considering the relatively minimal inferred weights associated with these three migration signals, it is posited that such patterns can be attributed to genetic introgression among *Oryza sativa* populations, as opposed to hybridization events. This interpretation underscores the subtle yet significant gene flow that characterizes the genetic landscape of these populations, distinguishing introgressive events from the more substantial genomic mergers typically associated with hybridization.

### Analysis of genomic mutation sites unveils differential positive selection across five *indica* super-hybrid rice varieties

Super-hybrid rice exhibits strong stress resistance and the potential for increased yield, underscore the importance of genomic features to comprehend the genetic traits of super-hybrid rice. In this study, we selected five *indica* super-hybrid rice varieties, namely Liangyoupei9 (LYP9), Y_liangyou_1 (Y1), Y_liangyou_2 (Y2), Y_liangyou_900 (Y900), Xiang Liangyou 900 (CY1000), each with different yield gradients. We also included the parental progenitors, Y58S, R900, Guanxiang_24S (GX24S), PA64SA, Yuanhui_2 (YH2), totaling 11 varieties. We obtained genomic NGS reads amounting to 478.86 M (71.67 GB bases) and proceeded to analyze their genomic mutation profiles (see methods). After filtering, our data exhibited a clean data rate of > 98% (approximately 470.86 M reads, 70.63 GB bases in total) with all samples achieving Q30 scores > 95% and mapping rates to the reference genome > 98%, indicate high data quality (Table S9). Through genomic variant analysis, a total of 26,715,592 SNP loci, 5,327,825 InDel loci, and 175,504 SVs loci were identified among five rice hybrids and their parental progenitors (Fig. 3, Table S10), with most sites exhibiting an allele frequency > 0.9, and minor genomic variation differences between samples (Fig. 3a, Fig. S5). We conducted a statistical analysis of the distribution of SNPs and SVs sites across genome regions, and the results indicate that the distribution trends across different samples’ genome regions are nearly consistent (Fig. S6, Fig. S7, Table S11). Notably, the ratio of missense to synonymous mutations (M/S) in each sample was > 1 but showed fluctuations (Fig. 3b), possibly indicating that *indica* populations are experiencing varying degrees of positive selection pressure, which may be closely related to differences in modern breeding goals for varieties. Additionally, we analyzed the shared mutation sites between the hybrids and progenitors for the five super-varieties, revealing that super-hybrid rice Y1 and Y900 shared more paternal genotypes, whereas the other three super-hybrid rice varieties shared slightly more maternal genotypes (Fig. 3c). Lastly, we mapped the high-quality reads from the above-cultivated rice NGS to the Nipponbare TE database and counted the relative mapping rates of reads in each type of TE to quantify the relative abundance of TE in each genome (see Methods). A comparative analysis through heatmap visualization of TEs abundance demonstrated a consistent pattern across various cultivated rice varieties. Notably, DNA transposons predominantly aggregated within the DNA/MuDR family. Conversely, RNA transposons exhibited a principal clustering within the Long Terminal Repeat (LTR) family, as illustrated in Supplementary Figure S8.

**Fig. 3.**
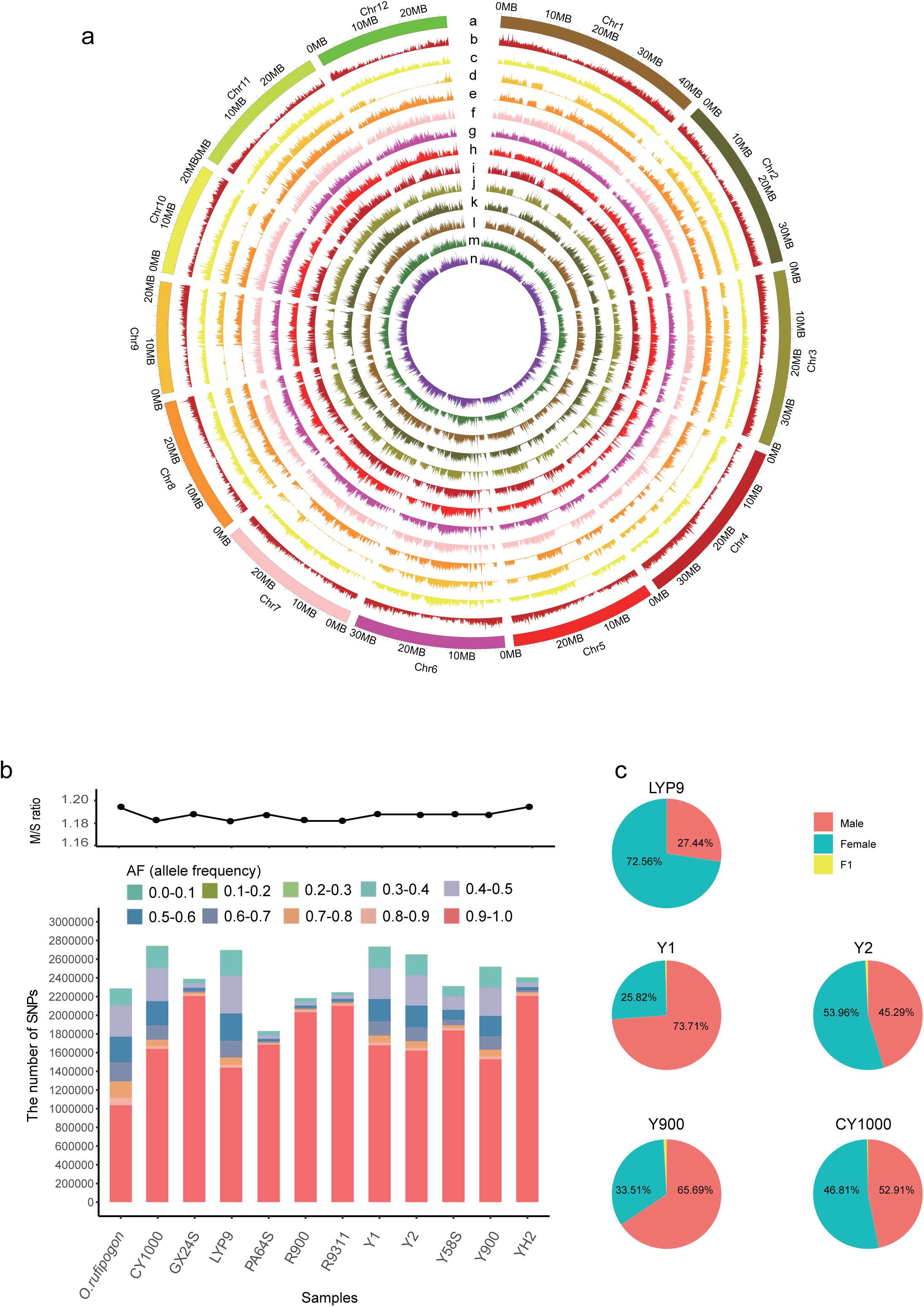
Genomic variants landscape of five *super-*hybrid rice and their hybrid progenitors. (a) Circos plot of SNP density within the genomes for the five super-hybrid and their hybrid progenitors, mapped in 1000 kb sliding windows. The outermost ring (a) represents 12 rice chromosomes, marked in Mb units. The second ring (b) illustrates gene density. Rings (c-n) represent SNP densities for rice verities: LYP9, PA64S, Y58S, R9311, Y1, Y2, YH2, R900, Y900, GX24S, CY1000 and *O. rufipogon*. (b) The top graph shows the ratio of non-synonymous to synonymous mutations, indicating the functional impact of genetic variation. The bottom bar chart displays the allele frequency (AF) distribution across different frequency spectra for the studied rice samples, revealing the polymorphism landscape. (c) Pie charts depict the genotype composition of the super-hybrid rice varieties in comparison to their parental progenitors, illustrating the inheritance pattern and genetic contribution from each parent.

### Genetic background analysis reveals *japonica* ancestry in the five *indica* super-hybrid rice varieties and frequent recent genetic exchange between the *indica* and *japonica* subgroups gives rise to diverse hybrid rice varieties

To delve deeper into the genetic history of *indica* super-hybrid rice, we performed genetic relationship analyses based on 90,113 high-quality SNPs in population samples from five super-hybrid rice and their progenitors. These were compared with *japonica* rice at the genomic level. HyDe analysis revealed that *indica* varieties inherited a degree of genetic material from *japonica* (Fig. 4a, Fig. S9). The above phenomenon could be attributed to two factors: 1) Inheritance from parental varieties such as R900, PA64S, Y58S, and R9311, which possess a *japonica* genetic background ^29,67^, and 2) the introduction of the *japonica* genetic background is attributed to complex gene exchange between the *indica* and *japonica* subgroups, as highlighted in previous studies ^42,68^. Observation of the heritability of super-hybrid rice and its parents revealed that parental genetic contributions to hybrids in LYP9, Y1, Y2, CY1000, and other super rice predominantly originated from the maternal side. Y900, however, was the more influenced by paternal genetics. The Heritability detection Z-score far exceeded |±3| and had a P-value of 0, indicating high credibility (Fig. 4a, b). The trend of parental genetic sharing in all varieties, except Y1, is consistent with the conclusion of genomic mutation sharing between offspring and parents in the previous section (Fig. 4b, Fig. 3c). The network genetic relationships constructed from the HyDe analysis (Table S12) among all *indica* varieties revealed intricate genetic relationships within *indica* subspecies (Fig. 4c,). In contrast, the admixture program ^69^ was used to analyze the genotype data of 90,113 SNPs, setting K from 3 to 15, to explore the genetic structure of these 11 *indica* rice varieties (Fig. 4d). By inferring the best K value, we identified two breaking points (K at 6 and 11). At K = 6, each super-hybrid rice possesses two ancestors with a mixing ratio of approximately 0.5, and the genetic composition of the super-hybrid rice is consistent with the parental selection in its breeding lineage. When K = 11, the cross-validation error is minimized, and since the population number is also 11, the estimated number of ancestors should be consistent with the population number. However, CY1000, R900, and LYP9 did not exhibit the characteristics of independent lineages, and the genetic composition of CY1000 included an unknown ancestor. We believe that this is more likely due to a bias caused by insufficient population numbers.

**Fig. 4.**
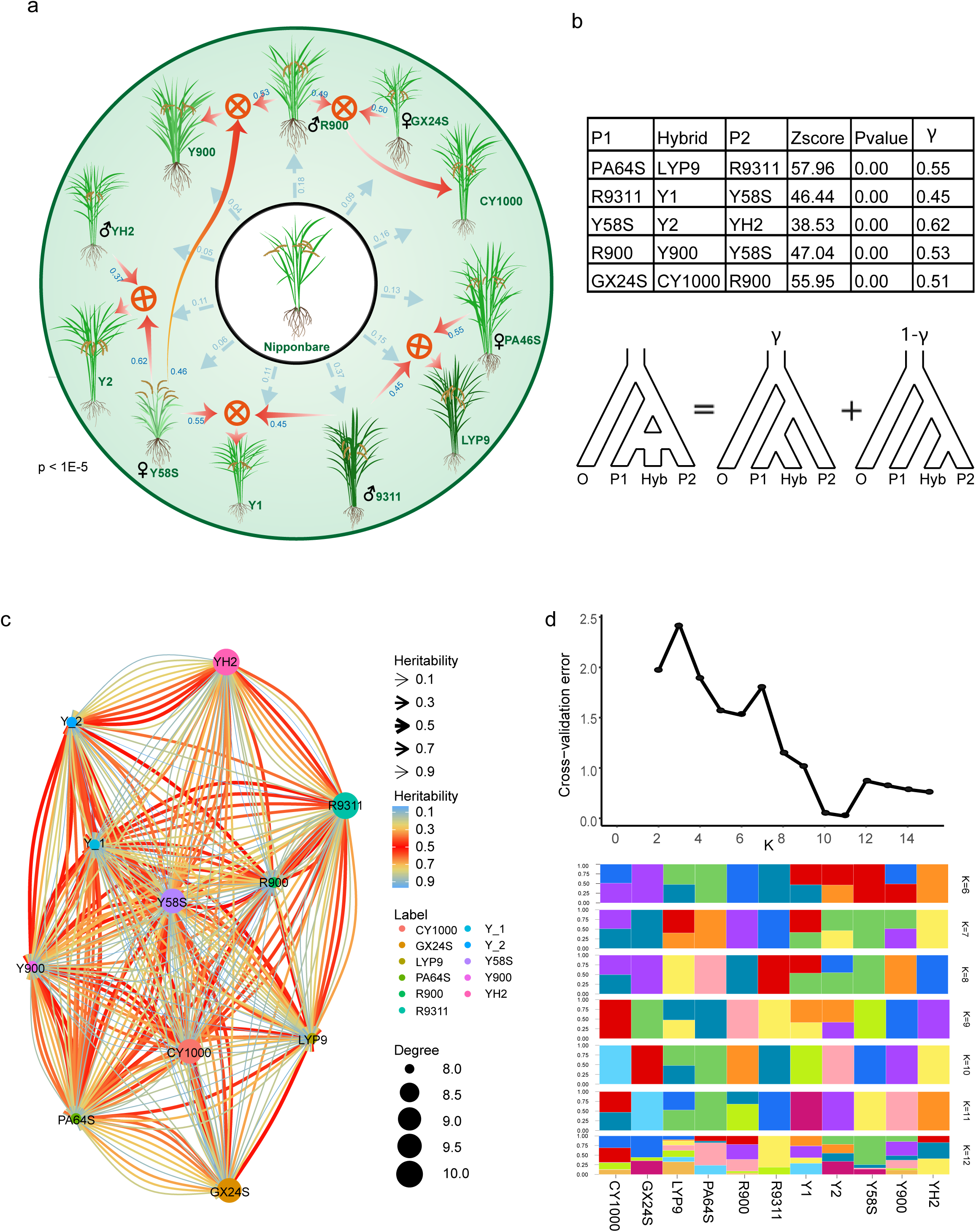
Analysis of hybridization history, population structure and genetic admixture in five elite super-hybrid rice varieties with their progenitors. (a) This circular diagram presents the hybridization history for each of five elite super-hybrid rice varieties as detected by HyDe, with significant hybridization events indicated where P-value < 0.05. The diagram highlights the complex interactions and genetic integration among the super-hybrids and their parental progenitors. (b) The table displays quantitative results from HyDe analysis, specifying the genetic proportions contributed by both parental progenitors to the super-hybrid *indica* rice varieties. The underneath accompanying schematic depicts the conceptual model for hybrid detection, representing hybrid (Hyb) as a composite of genetic input from two parental progenitors P1 (γ) and P2 (1-γ), reflecting their respective heritability contributions. (c) The network graph illustrates the genetic heritability resulting from hybridization events among various rice varieties. Node sizes correspond to the degree of connectivity, while edge colors denote levels of heritability. Arrows point towards the hybrid offspring, indicating the direction of genetic contribution from the parents, with specific heritability values labeled alongside. (d) The top graph provides an estimation of the cross-validation error for determining the optimal number of ancestral populations (K), while the bottom admixture plot, categorized by the same optimal K, reveals the structural composition of genetic elements within each sample. The color coding represents unique genetic components derived from the ancestral populations.

In summary, the heritability and population structure analysis between hybrid rice and its parents vividly elucidate their complex hybridization history, consistent with the breeding history documented by breeders. Moreover, the recent signals of reciprocal hybridization between *indica* and *japonica* rice varieties explain the genetic factors underlying the formation of numerous elite hybrid varieties. This underscores the importance of deciphering more genetic resources in rice to guide the improvement of cultivated rice varieties.

### Differential expression patterns among various super-hybrid varieties with their progenitors and the prevalence of non-additive inheritance is likely to be major contributors to heterosis

Super-hybrid rice, renowned for its exceptional traits such as stress resistance and high yield, is pivotal to bolstering food security. Understanding the underpinnings of its superiority is therefore crucial. To delve deeper into the formation of hybrid advantages in super-hybrid rice, we analyzed the transcriptome data of samples including LYP9, Y900, and CY1000 with their progenitors (Table S13). We conducted pairwise comparisons for each super-hybrid rice variety with their parental progenitors, retaining genes with significant expression differences in any comparison group. The clustering of all differentially expressed genes unveiled a distinct tissue-specific trend across all samples (Fig. 5a, Table S15). To enhance the observation of expression patterns between offspring and parents, we further employed Mfuzz ^70^ to cluster all genes based on their expression profiles across progenitors and offspring. This resulted in six clusters (Figure 5b, Table S16). Notably, clusters 3, 4, and 5 exhibited gene expression of hybrid offspring higher than one or both parents, suggesting that these highly expressed genes in offspring may contribute to trait superiority. Additionally, we classified genes based on their expression relationships between offspring and parents in different tissue samples of the LYP9, Y900, and CY1000 hybridization combinations (see Methods). The findings indicate that additive genes (A) account for approximately 25% in all offspring tissue samples of these three super-hybrid rice combinations, with non-additive genes predominating (Figure 5c, Table S17). A previous study on the LYP9 super-rice hybrid combination indicated that the predominance of non-additive genes may be a key factor in the formation of the trait dominance ^40^. Our study not only incorporates data from the LYP9 with its progenitors but also includes samples from the recently released Y900 and CY1000 super-hybrid rice varieties with their progenitors. The similar results observed in these super-hybrid rice varieties further underscore the significant role of non-additive expression genes in the formation of hybrid advantages in super rice.

**Fig. 5.**
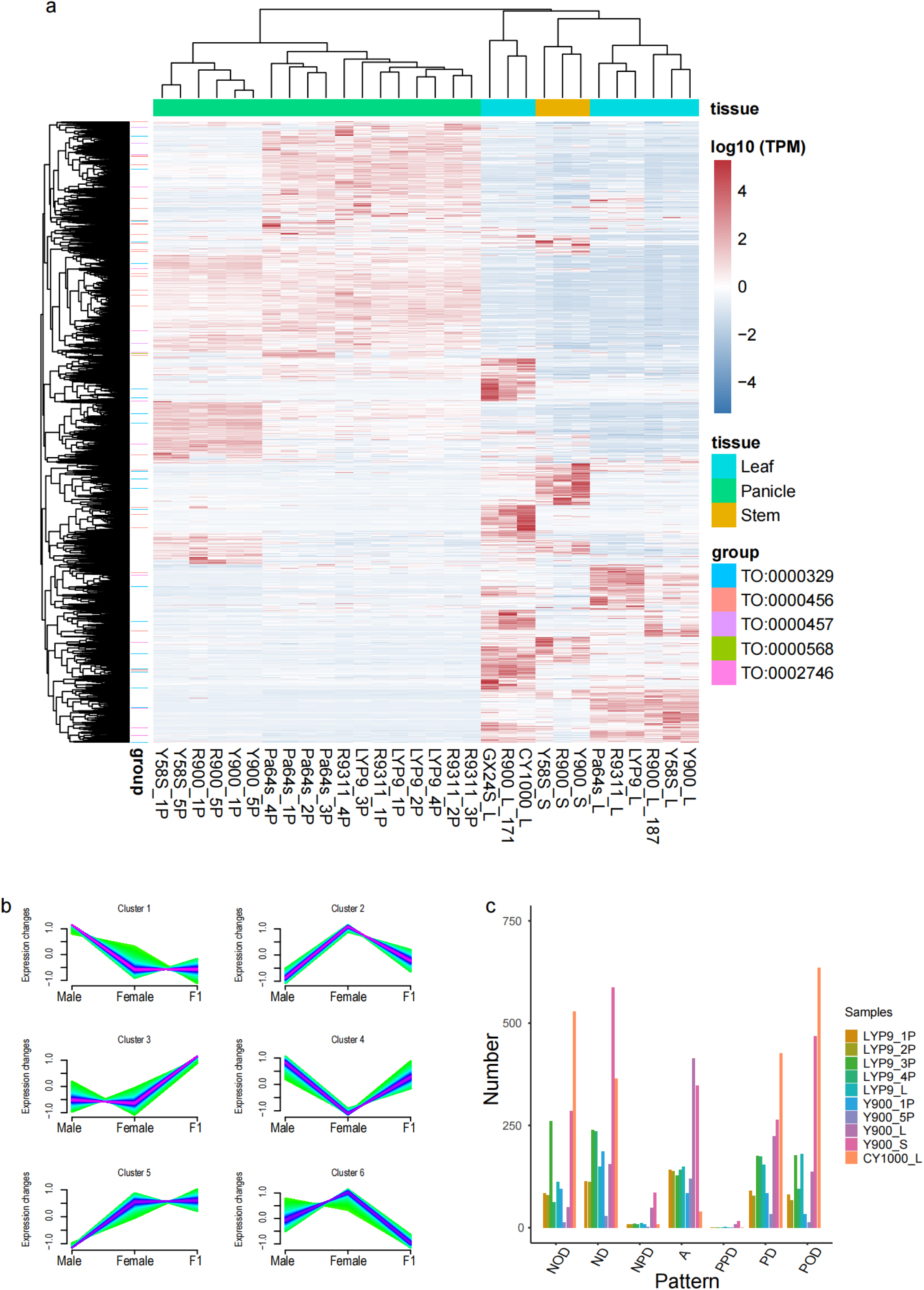
Gene expression profiling in elite super-hybrid rice varieties with their parental progenitors. (a) Heatmap illustrating the patterns of differentially expressed genes across various tissues in three super-hybrid rice varieties (LYP9, Y900, and CY1000) and their parental progenitors. The horizontal color bands represent different tissue types with corresponding expression transcripts per million (TPM) value, while vertical color bands classify genes according to Trait Ontology (TO) categories, sourced from the China Rice Data Center. Expression levels are color-coded, with red indicating higher expression and blue indicating lower expression. (b) Line graphs depict the general expression trends across all differentially expressed genes within the three super-hybrid rice varieties and their parental lines, with separate trends plotted for male, female, and F1 hybrid tissues. (c) Bar graph categorizing the number of differentially expressed genes across the super-hybrid varieties, with color coding representing distinct patterns of gene expression in response to various biological and experimental conditions (‘A’ indicates additive gene, all others are non-additive).

### eQTL analysis

The influence of genomic mutations on gene regulation has been the subject of extensive research, with numerous studies underscoring their substantial impact ^45,71,72^. Utilizing the FastQTL software, we correlated transcriptome expression data with genome mutation data for LYP9, Y900, and CY1000. We identified 82, 370, and 205 genes (termed eQTL genes) with high signal values in LYP9, Y900, and CY1000, respectively (LYP9: P-value < 1e^-4^; Y900, CY1000: P-value < 1e^-5^) (Table S18). These genes included 25 known phenotype unique genes primarily associated with biotic and abiotic stresses, plant height, roots, stems, leaves, spikelet, and fruit morphology phenotypes (Fig. 6, Table S19). For example, the *OsRBD1* gene can enhance yeast tolerance to abiotic stresses such as salt and osmotic stress by interacting with *OsSRO1a* gene ^73^. In our study, the *OsRBD1* gene is a highly associated locus in Y900 and CY1000, especially in CY1000, a well-known saline-grown rice ^74^, this suggests the *OsRBD1* gene may plays a crucial role in salt tolerance. In comparison to the wild-type the *LFS* mutant showed a decrease in plant height and spike length, grain number per spike, 1,000-grain weight and spikelet fertility ^72^. Additionally, only 31 overlapping genes were identified between Y900 and CY1000 among these eQTL genes, with the remainder being specifically expressed in their respective samples (Fig. S10).

**Fig. 6.**
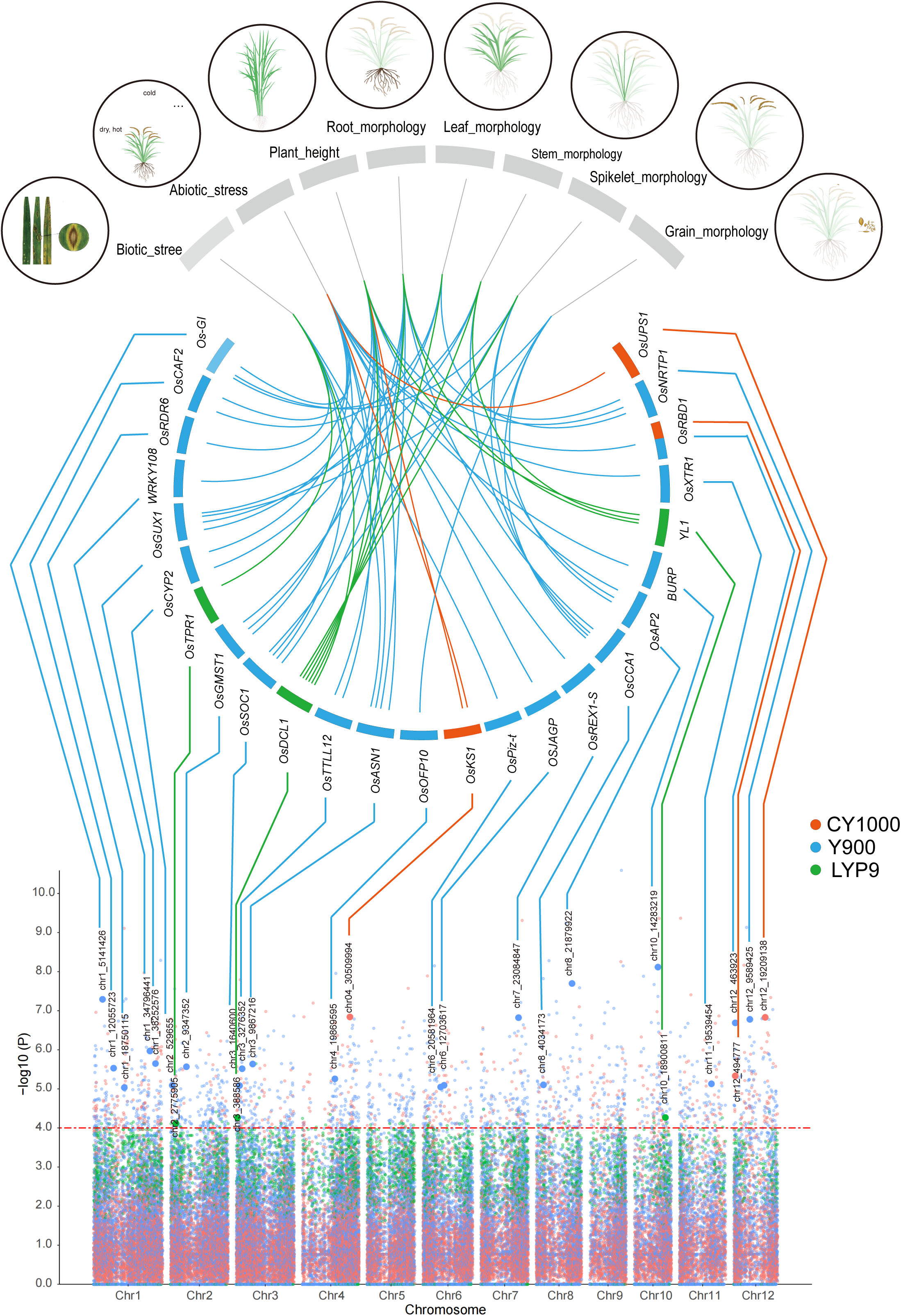
eQTL mapping of morphological traits in three elite super-hybrid rice varieties. Top panel: the Circos plot illustrates the association between morphological traits and linked candidate genes within significant SNP loci for the rice varieties CY1000, Y900, and LYP9. Traits such as plant height, root, leaf, stem, spikelet, and grain morphology are depicted in the outer ring with corresponding candidate gene connections drawn to the inner SNP locus ring. Bottom panel: A Manhattan plot presents a comprehensive view of SNP loci associated with the candidate genes for morphological traits across all rice chromosomes. Each dot represents an SNP, color-coded to distinguish the contributions from each of the elite super-hybrid rice varieties CY1000 (blue), Y900 (green), and LYP9 (red).

To further investigate the potential molecular functions of eQTL loci across different hybridization combinations, we performed GO/KEGG enrichment analysis for the aforementioned eQTL genes. In GO enrichment analysis, the LYP9 hybridization combination group was mainly enriched to 31 GO terms (Fig. S11, Table S20), including 14 Molecular Functions (MF) and 17 Biological Processes (BP), predominantly concentrated in DNA-templated transcription (GO:0006351). The Y900 group was annotated to 99 GO categories (Fig. S11, Table S21), including 51 BP, 19 Cellular Components (CC), and 28 MF, primarily annotated on stress, defense responses to external (GO:0050896, GO:0006950, GO:0006952). CY1000 mainly included 36 BP, 14 CC, and 15 MF, predominantly enriched in metabolic processes (Fig. S11, Table S22). KEGG annotated 17 significant unique pathways, with LYP9, Y900, and CY1000 belonging to three, six, and ten, respectively (Fig. S12, Table S23). Notably, Y900 and CY1000 shared pathways for homologous recombination (ko03440) and nucleotide excision repair (ko03420) pathways.

### De novo SNP-associated genes are likely contributed to yield advantage formation in super-hybrid rice varieties

Through examination of genomic mutation sites, we postulated that de novo-originated SNP sites (specific to offspring) may have significantly contributed to the genetic underpinnings of yield formation in high-yielding super rice. In our analysis of five hybrid-rice combinations, we identified a total of 496 de novo origin sites (109 in Y1, 80 in Y2, 268 in Y900, 39 in CY1000), whereas no de novo origin SNP sites in offspring were found in the LYP9 analysis. Through annotation, we identified 103 de novo sites involving 81 genes (Table S24), including 11 markers associated with biotic and abiotic resistance, plant morphology, and other phenotypic traits (Fig. 7a, Table S25). When correlating these 81 genes with the transcriptome data, only 21 genes were expressed in at least one sample of the three hybrid-rice combinations described above, including six marker genes (*Os03g0191000*, *Os04g0121100*, *Os11g0134700*, *Os10g0561400*, *Os03g0181500*, and *Os03g0758900*), showing varying expression trends across different hybrid rice combinations (Fig. 7b, c). Specifically, the heat tolerance gene *Os03g0191000* exhibited lower expression in CY1000 compared to its parents ^75^. *Os11g0134700*, which exhibits diverse expression patterns under various biotic and abiotic stress conditions, such as drought, salinity, cold, and heat, and biotic stresses (including infection by *Magnaporthe Oryzae*, and *Xanthomonas Oryzae* pv. *Oryzae* and *Rhizoctonia solani* nematodes) ^76^ had consistently lower expression levels than those of the parents in both Y900 and CY1000. The overexpression of *Os10g0561400* may contribute to increased cold tolerance in Y900 ^77^. The gene *Os03g0181500* influences post-germination plant growth, with higher expression in LYP9 and CY1000 than in either parent ^78^. The gene *Os03g0758900*, which positively regulates the expression of disease-resistance genes, had an expression level higher than that of at least one parent in all three super rice combinations ^79^. These genes may play a key role in the development of superior traits in super rice.

**Fig. 7.**
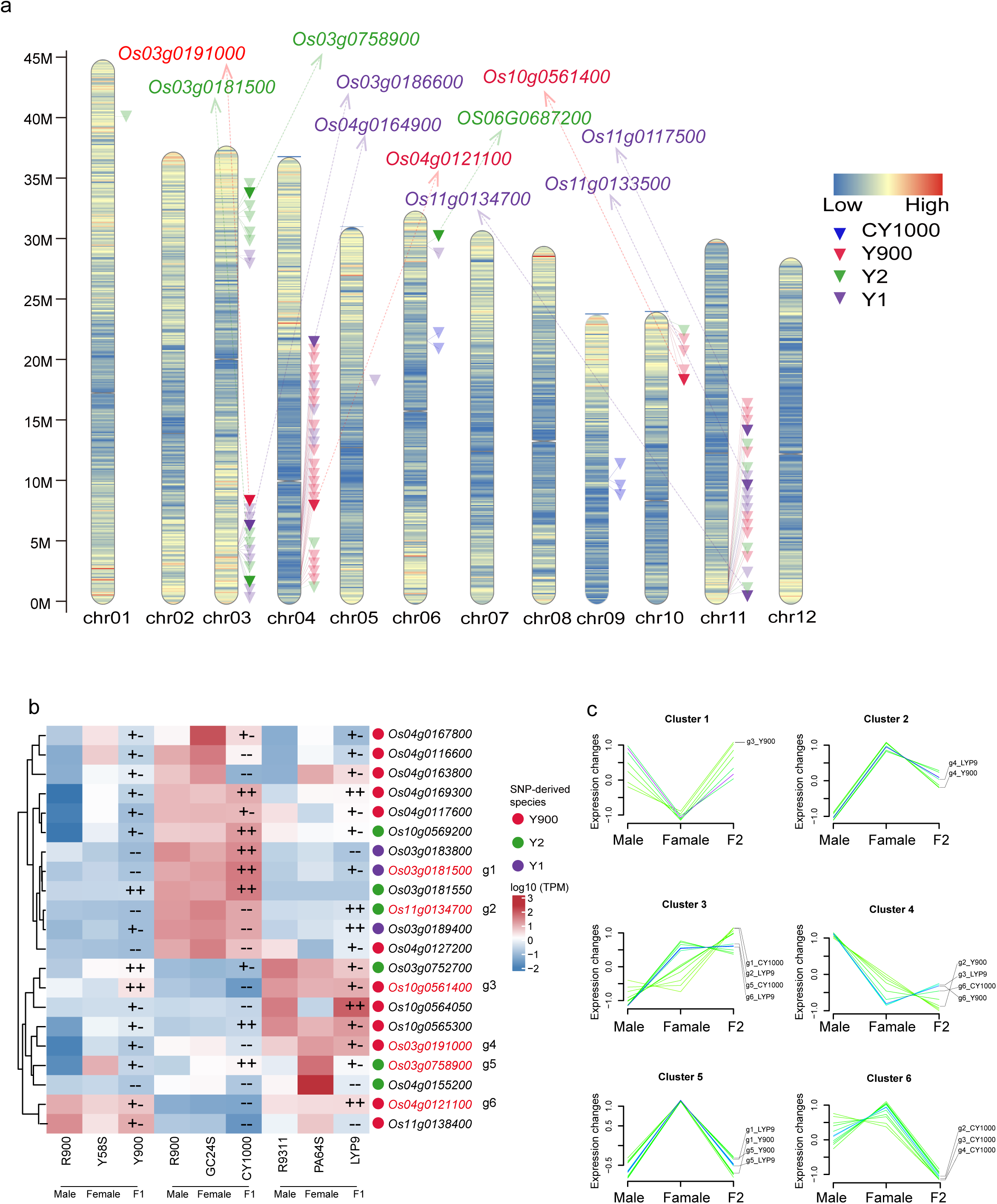
Distribution and expression pattern of genes with de novo SNPs identified for four elite super-hybrid rice varieties. (a) The graphic displays the density of genes across the 12 chromosomes of rice, with the color scale indicating gene density. Inverted triangles pinpoint the locations of genes with identified de novo SNP loci, and genes with functional descriptions are labeled above each chromosome. (b) The heatmap details the differential expression levels of the genes identified in part (a) with genes with functional descriptions highlighted in red text. The colored scale on the right indicates the gene expression level, while the colored circle symbols next to gene names for super-hybrid rice varieties. The legend defines the symbols: ’++’ indicates higher expression than both parents, ’+-’ indicates higher expression than one parent, and ’--’ indicates lower expression than both parents. (c) The line charts represent the expression profiles of genes with functional descriptions identified in part (b) across male, female, and F2 hybrid samples from the super-hybrid rice combinations, with line colors corresponding to the gene come from different super-hybrid rice varieties.

## Discussion

The origins of Asian rice domestication remain contentious, with prevailing hypothesis divided between a single domestication event and multiple independent domestications. Substantial research supports both hypotheses. Studies favoring a single origin tend to focus on specific genetic loci, whereas those advocating for multiple independent origins emphasize whole-genome variation. However, the intricate evolutionary history of Asian rice, further complicated by recent decades of breeding efforts ^11^, has resulted in complex genetic landscape among its subspecies. This complexity can significantly influence the accuracy of the subspecies origin hypotheses based on a limited number of genetic loci ^80–82^. The recent availability of high-quality genomes of *Oryza* spp. facilitates the further elucidation of phylogenetic relationships within cultivated rices. Our study examined the complete gene set of 34 genomes, summarizing 39,984 multi-copy gene trees to construct a robust *Oryza* phylogeny. This approach minimized distortions caused by small gene scales and gene flow between certain subspecies, thereby circumventing misrepresentation of species relationships ^83^. Consequently, our research contributes to the advancement of resolving the origins of Asian rice domestication. Our gene-level analysis across a broad range of *Oryza* spp. provides comprehensive evidence of the origins and domestications in Asian rice. Utilizing the species tree summarized by multi-copy gene family trees, we discovered that *indica* and *japonica* rice formed independent lineages with distinct *O. rufipogon* species as sister group, indicating separate origins from different *O. rufipogon* progenitors. Moreover, at the most common ancestor of the *indica* and *japonica* subgroups, the divergence time showed that *japonica* > *indica*, which may be due to incomplete sampling of wild species in the species tree. Previous studies have established that Asian wild rice is divided into several verities, and *O. nivara* also originates from *O. rufipongon* ^19,84^. Therefore, we speculate that there exists an unknown Asian wild rice ancestral species at the most common ancestor node of the *aus* and *indica* clades, so the actual divergence time (0.25 of this node is likely similar to that of the *japonica* rice in our study. In addition, the trend in genome-wide syntenic gene pair Ks distribution further corroborates that the age of two major subspecies origin is close to each other ^10,20,22,85^. Concurrently, analyses of genes within the nucleotide low-diversity region revealed parallel domestication times for the *indica* and *japonica* subspecies, thereby confirming the conclusion of independent domestication.

A previous study has suggested that *O. nivara* is a wild ancestor of the *aus* group ^12^. By incorporating a more comprehensive array of wild rice species, we identified a close relationship between *aus* and the annual wild rice *O. nivara*, whereas *indica* was closer to the wild rice *O. rufipogon* w1654, with the *aus* and *indica* clades being sister groups. This evidence supports the independent origins of *aus*, *indica*, and *japonica* from different wild rice varieties, which is consistent with recent studies that have posited *O. nivara* as a separate lineage ^10,80,86^. Notably, although previous studies suggested that *O. nivara* originated multiple times from different *O. rufipogon* populations ^84,86^, our data describe *O. nivara* and *O. rufipogon* w1654 as members of the sister taxon. (Fig. 1a). This may be due to under-sampling of our wild rice genome. Therefore, we hypothesized that *aus* and *indica* rice were differentiated from this unknown type of Asian wild rice, *O. rufipogon*.

Despite preponderance evidence supporting the polyphyletic origin of Asian rice, gene flow between Asian cultivated rice subspecies has also been observed, particularly in the introgressive influence of *japonica* on *indica* and its ancestors ^20,42,63^. In our study, extensive recent genetic introgression from the *japonica* subgroup to the *indica* subgroup was observed (Fig.2, Fig. 4, and Fig. S9), which is closely linked to modern breeding practices. Furthermore, the analysis of duplicate genes at the ancestral node of Asian rice (genes conforming to the ABAB pattern, indicating no hybridization between the MRCA of rice) further bolstered the hypothesis of independent origins of the *indica* and *japonica* subgroups. However, we noted a few gene duplications occurred in the MRCA of rice where genes were missing from either the *indica* or *japonica* branches, which we speculate may be due to the following: 1. different evolutionary patterns resulting in gene loss from certain subpopulations. 2. insufficient species coverage leading to undetected genes in certain subpopulations. 3. few gene flow events between subpopulations in the MRCA of *O. sativa*, especially from *japonica* to *indica* or *aus*, resulting in the absence of the gene in a particular branch of *indica* or *aus*. A prior study indicated that Asian wild rice is a hybrid swarm with extensive gene flow and feralization from domesticated rice, gene flow from domesticated rice to wild rice populations could result in genetic similarities between them ^81^, as observed at the *indica* rice ancestral node (Fig. S2 d, f), where a large number of wild rice share some genetic background with cultivated rice that may also include gene flow from cultivated to wild rice. In addition, genetic introgression among wild rice is not limited to the African AA genome type ^64^, but also occurs among other types of wild rice genomes. These finding provides a more comprehensive explanation of the genetic background of Asian rice and contributes to the understanding of its diversity and evolutionary dynamics.

The continuous development of breeding technologies has led to significant improvements in super-hybrid rice. Research has shown an upward trend in yield for five super-hybrid rice combinations, LYP9, Y1, Y2, Y900, and CY1000, with CY1K reached 18.0 t/hm2 ^29,87^. The successful breeding of these high-yielding varieties makes it possible to analyze their underlying genetic mechanisms. Genomic data analysis of these five hybrid combinations provides a comprehensive understanding of the genetic relationships and genomic characteristics of the progeny and parents. Allelic frequencies were generally high across different samples, with slight variations in the density distribution. These characteristic differences may be associated with positive selection acting on the relevant genomic regions of the rice varieties. Moreover, previous transcriptomic studies have suggested that non-additive effects inherently refer to the functional underperformance of two homozygous alleles from parents with insufficient background, whereas in the same background, one allele in the heterozygous F_1_ may fully function, exhibiting partial dominance, dominance, or even over-dominance ^40^. Our findings that the transcriptomes of the super-hybrid rice combinations were distinctly tissue-specific and that a large number of alleles were expressed at higher levels in the hybrid progeny than in one or both of the parents, with a predominance of non-additive genes, led us to infer that they were associated with heterosis. This results in the expression of traits that are superior to those of the parents after crossing. Therefore, we hypothesize that this part of the gene may also play an important role in the formation of heterosis.

Numerous studies have confirmed the critical effect of genomic mutations on gene expression ^45,71,72^. It has been demonstrated that phenotypic traits, such as hair color, shape, and thickness, are regulated by certain associated loci ^44^. In our eQTL analysis, a group of stress-related genes significantly associated with mutant loci were identified. In addition, we found a large number of enriched GO terms in LYP9 related to environmental stress and defense, which illustrates the important role of these eQTL genes for LYP9 resistance traits. Furthermore, shared nucleotide excision repair and homologous recombination pathways in Y900 and CY1000 were observed in KEGG enrichment analysis. Previous research has indicated that DNA damage arises from both endogenous and exogenous factors ^88^. Nucleotide excision repair is the primary pathway for removing DNA damage ^89^, and homologous recombination plays a crucial role in DNA repair and interchain crosslinking (ICL) ^90^. These pathways were significantly enriched in Y900 and CY1000, whereas no relevant pathways were identified in LYP9. Therefore, we hypothesized that these factors above were closely associated with the resistance traits of Y900 and CY1000.

## Conclusion

This study was designed to shed light on the origins of cultivated rice and the genomic characteristics underlying that underpin the heterosis of super hybrid rice yield traits by integrating multi-omics data. By amalgamating the complete genomes of 33 high-quality Oryzeae species along with one outgroup *Brachypodium distachyon* genome, we constructed a robust phylogenetic relationship by summarizing 39,984 multi-copy gene trees. Our analysis revealed that *indica* and *japonica* as distinct lineages, supporting their independent origins from different wild rice species. Through assessing the collinear gene pairs and Ks distribution between cultivated rice subspecies and closed wild rice varieties, as well as estimating the divergence time and Ks distribution of 54 putative domestication genes identified from nucleotide low-diversity regions, we found that although the overall domestication time of *indica* and *japonica* subspecies is similar. However, Ks analysis based on 11 key domestication genes revealed that the domestication time of some key genes related to morphology and grain shape in *indica* precedes that of *japonica* rice suggesting that the domestication of cultivated rice subspecies might have occurred in stages, with early domestication of certain traits in *japonica* rice, followed by simultaneous large-scale domestication of both *indica* and *japonica* rice to form modern subspecies. Analysis of the topological pattern of 1,383 gene duplication in the MRCA of *O. sativa* did not support recent hybridizations between the MRCA of *indica* and *japonica* subspecies, further supporting their independent origins. Hybridization analyses, employing datasets of 1,300 single-copy orthologous genes alongside 669 orthologous genes originating from gene duplication events in the MRCA of *Oryza sativa*, corroborate the hypothesis of ancient introgression from early divergent Oryzeae species contributing to rice speciation. This evidence, although limited, suggests a nuanced role of hybridization in the evolutionary origins of the MRCAs of *indica*, *japonica*, and *aus* subspecies. Genetic network analysis of whole-genome sequencing data from wild rice and cultivated rice subspecies, as well as analysis of hybridization, genetic networks, and population structure of whole-genome NGS data from five super rice combinations, revealed hybridization signals between *indica* and *japonica* rice and among *indica* rice subspecies, highlighting the complex breeding history of cultivated rice and the importance of introduction of cultivated rice subspecies for the breeding of elite varieties.

On the other hand, we conducted whole-genome NGS sequencing of five major hybrid *indica* rice varieties (LYP9, Y1, Y2, Y900, and CY1000) and their progenitors. Mutation analysis revealed genomic variation characteristics and different degrees of positive selection on those multiple hybrid-rice varieties. Hybridization detection and population genetics analysis strongly demonstrated extensive hybridization and introgression between the five commercialized super hybrid varieties and *indica* rice, indicating the integration of multiple genetic resources during the cultivation of super hybrid rice. Transcriptomic RNA-seq data analysis of three super hybrid varieties (CY1000, Y900, and LYP9) and their parental progenitors highlighted different gene expression patterns between hybrid varieties, emphasizing the potential importance of non-additive genes in trait superiority. eQTL analysis, integrating gene expression data with mutation maps, revealed that the resistance mechanisms of Y900 and CY1000 might be closely associated with shared recombination and DNA repair pathways. Additionally, it uncovered a set of genes that may influence the formation of advantageous traits.

In conclude, our study provides comprehensive evidence to support the independent domestication hypothesis for the origins of the *indica* and *japonica* rice subspecies and their parallel domestication histories. Through analyses at different levels focusing on ancestral nodes of *indica* and *japonica* subspecies, we elucidate the potential impact of ancient genetic lineages on the formation of cultivated rice and the complex gene flow history among cultivated rice varieties. Additionally, analyses at the genomic and transcriptomic levels of super-hybrid rice provide new insights into the genetic basis of yield advantage and release a set of genes associated with superiority, thus providing data support for broadening breeding resources.

## Materials and Methods

### Experimental materials and data sources

The dataset for this investigation comprised 34 genomic coding sequences, sourced from eminent public repositories: the Rice Genome Hub (https://rice-genome-hub.southgreen.fr), the National Genomics Data Center (https://ngdc.cncb.ac.cn/), and the National Center for Biotechnology Information (https://www.ncbi.nlm.nih.gov) (refer to Table S1). For the construction of interspecific genetic networks, whole-genome sequencing datasets were retrieved from the European Nucleotide Archive (ENA, https://www.ebi.ac.uk/ena/browser/home), as documented in Table S8, employing TreeMix v1.13 software ^65^ for interspecific genetic network analyses. Transcriptomic RNA-seq datasets includes tissue samples from roots, stems, and leaves of three elite super-hybrid rice varieties including Liangyoupei9 (LYP9), Y_liangyou_900 (Y900), and Xiang Liangyou 900 (CY1000) alongside their progenitor strains were curated from the NCBI Sequence Read Archive (https://www.ncbi.nlm.nih.gov/sra) and the CNCB database (https://ngdc.cncb.ac.cn/), as detailed in Table S13. Five super-hybrid rice varieties including Liangyoupei9 (LYP9), Y_liangyou_1 (Y1), Y_liangyou_2 (Y2), Y_liangyou_900 (Y900), Xiang Liangyou 900 (CY1000) were planted in Erlang Village, Jiusuo Town, Pingqiao District, Xinyang, Henan (114.08°E, 32.13°N), while their respective parents including Y58S, R900, Guanxiang_24S (GX24S), PA64SA, Yuanhui_2 (YH2) of the super-hybrid rice were cultivated in Jiaojiao Village, Jiusuo Town, Ledong County, Hainan Province (109.17°E, 18.73°N). Post-harvest, young foliar specimens were promptly submerged in liquid nitrogen for preservation and subsequently stored at -80°C. Genomic DNA was extracted, and high-throughput sequencing libraries were synthesized utilizing the Nextera DNA Flex Library Prep Kit (Illumina, San Diego, CA, USA). Subsequently, paired-end 150 sequencing is performed on the Illumina NovaSeq 6000 platform. In total, approximately 71.67 GB of bases was obtained.

### Genome-scale phylogenetic analysis with cultivated and wild rices

We curated a dataset comprising 33 high-quality genomes within the Oryzeae and *Brachypodium distachyon* serving as an outgroup, to refine the phylogenic resolution in subgroups level (Table S1). The incorporation of coding sequences derived from the aforementioned genomes effectively minimize errors attributable to the disparate evolutionary trajectories of partial genes, which arise from varying degrees of bottleneck effects. The above approach also circumvents inconsistencies in outcomes that may result from partial gene flow between ancestral populations of cultivated rice originating from distinct sampling locations, as well as between cultivated rice and its ancestral groups. All annotated protein sequences from a total of 34 genomes were clustered into 39,984 orthologous groups (OGs) using OrthoFinder v2.5.5 ^91^ with default parameters. Protein sequences corresponding to echo OGs were extracted, and MAFFT v7.508 with parameter “--auto” ^92^ was performed for multiple sequence alignment. The protein alignments were converted into nucleotide multiple sequence sequences, guided by the original nucleotide coding sequences (CDS) by PAL2NAL ^93^. The maximum likelihood gene family trees were inferred using IQ-TREE v2.2.0.3 under the GTR+G model with 1000 bootstrap replicates ^94^. Finally, all multi-copy gene family trees were summarized into a coalescent species tree using ASTRAL-Pro 2 (v1.13.1.3) ^83^ with default parameters.

### Molecular clock Analysis

Molecular clock was used to estimate the divergence time within the Oryzoideae. On one hand, we filtered out genes with lengths smaller than 1,000 bp from the 1,300 single-copy gene clusters, resulting in 225 single-copy gene clusters used for species divergence time analysis. On the other hand, genes from 15 regions of low nucleotide diversity reported in a previous study ^10^ and genes related to yield collected from the China Rice Data Center (https://www.ricedata.cn/) were intersected; then, this set was intersected with 1,300 single-copy gene clusters obtained from species phylogenomic analysis; finally, genes shorter than 1,000 bp were filtered out, resulting in 54 clusters of orthologous single-copy genes of putative domestication genes used to assess the domestication time among different subspecies. We extracted respectively the protein sequences of all the genes of these two gene clusters set using custom Perl scripts and then aligned using MAFFT v7.508 ^92^ for multiple sequence alignment. Subsequently, PAL2NAL ^93^ was utilized to convert the alignments into CDS sequences. We concatenated the CDS sequences of the 225 gene clusters dataset and the 54-domestication gene dataset into separate sequence matrices, respectively. Then, we used MCMCtree ^95^ to infer the divergence time between species and the domestication time. Furthermore, to minimize potential systematic errors caused by multi-gene concatenation, MCMCtree was used to individually analyze the 54 putative domestication genes, and statistical analysis was conducted on the distribution of divergence node times to assess relative domestication timing.

### Ks analysis for syntenic and orthologous gene pairs to the divergence time of *indica* and *japonica* subspecies

The synonymous substitution rate (Ks) is typically not affected by selection pressure and can characterize the evolutionary history of species. First, we separately counted the Ks values of syntenic gene pairs between the Asian cultivated rice genomes and their corresponding MRCA species within their subgroups to assess the relative divergence time between species. Then, we assessed the genome duplication status by computing the Ks values of homologous gene pairs within the cultivated rice species itself. Homologous gene pairs were identified using Diamond v2.0.15.153 with parameters “-e 1e^-10^ --max-target-seqs 5 --masking 0 --no-self-hits --outfmt 6” ^96^. Syntenic gene pairs were identified and analyzed using MCscanX ^97^ with parameters “-k 100 -s 5 -e 1e^-10^”. The protein sequences of collinear gene pairs were aligned by MAFFT v7.508 with parameter “--auto” ^92^. The protein alignments were converted into nucleotide alignments by PAL2NAL ^93^. Ks values were calculated for each aligned syntenic gene pairs using kaks_Calculator v2.0 with model NG ^98^. On the other hand, *aus*_n22, R9311 and Nipponbare, which have better annotation quality, were used as representative species of *aus*, *indica* and *japonica* subspecies. For the 54 domestication gene clusters inferred from the previous molecular clock analyses in the regions of low nucleotide diversity, the genes of the representative species of subspecies *aus*, *indica* and *japonica* in the gene clusters were used as orthologous gene pairs with the corresponding genes of the MRCA wild species of subspecies *aus*, *indica* and *japonica*, and Ks analyses were carried out to assess the relative domestication times of the *aus*, *indica,* and *japonica* subspecies. In addition, for the more important 11 domesticated genes in rice ^42,53–55^, Ks values were calculated between species within the *indica* and *japonica* subgroups, respectively, and the MRCA species of the subspecies to which they belong. All the above analyses were graphed using the R package ggplot2 ^99^.

### Phylogenomic profiling for detection of gene duplication events

Gene duplication events at each node of the species tree were identified by using Tree2GD v2.5 with parameters “--bp=70 --sub_bp=70 --species=2 --root=MAX_MIX” ^47^. Using the rooted species tree as a framework, all the gene family trees were rooted and mapping onto the species tree for tree reconciliation analysis, resulting in the identification of 1,383 instances of gene duplication events. Subsequently, our custom Perl script was utilized to quantify the topological configurations of the gene trees implicated in these duplication events. A total of three topologies (ABAB, ABAx, and ABxB) of cultivated rice ancestor nodes were involved. The ABAB-type topologies suggest that subpopulation A and subpopulation B diverged independently at the ancestral node, whereas other types of topologies indicate potential gene flow between the two subpopulations.

### Identification of tandem gene duplication event

To investigate the origins of gene duplication events and determine whether such duplications in the ancestors of the genus *Oryza* were attributed to an allopolyploidy event, a comprehensive analysis was undertaken utilizing R9311 (representative of the *indica* subspecies) and Nipponbare (representative of the *japonica* subspecies) as model organisms. This analysis focused on the quantification of tandem gene duplications, defined by the spatial proximity of duplicated gene pairs. Specifically, a criterion was employed wherein a maximum of ten intervening genes between each pair was permissible, thereby facilitating a detailed examination of the gene duplication landscape within these genomes.

### TE annotation and association analysis for duplicated genes and the presence of TEs

To elucidate the origins of gene duplication events and assess the hypothesis that the extensive gene duplications observed in the ancestors of the genus *Oryza* were not derived by allopolyploidization, we conducted a comprehensive TE annotation and association analysis. Specifically, the genomes of the *indica* and *japonica* rice representatives, Nipponbare and R9311 respectively, were subjected to TE identification using the RepeatMasker v4.1.2 (https://www.repeatmasker.org/). The analysis employed parameters “-species rice -xsmall -s - norna -no_is -nolow -gff” and integrated the Dfam 3.8 ^100^ and Repbase databases (https://www.girinst.org/) for TE annotation. To explore the hypothesis that the observed extensive gene duplications may be mediated by TEs, we conducted a statistical analysis of TEs located within the 2 kb upstream and downstream regions of repetitive genes and other genes excluding repetitive genes, based on the genomic annotation of both Nipponbare and R9311. We utilized the chi-square test to statistically analyze the association between the presence of TEs and gene duplications, aiming to gain deeper insights into the role of TEs in the genome evolution of *Oryza* species.

### Identification and filtration of SNPs, structural variations (SVs) and of TE abundance by utilizing whole-genome sequencing data derived from super-hybrid rice and its parental progenitors

Genomic NGS data generated for the five super rice combinations in this study were used for genomic variant site analysis. The specific steps are as follows: Firstly, the raw data were filtered and quality controlled using Trimmomatic (v0.39, options: LEADING:10 TRAILING:10 SLIDINGWINDOW:4:20 MINLEN:36) ^101^ to obtain high-quality reads and high-quality reads were aligned to the Nipponbare genome using BWA software (0.7.1) ^102^ mem module to generate SAM format files. SAM files were sorted into BAM files using SAMtools v1.16.1 ^103^ and duplicate sequences were marked using the Picard software MarkDuplicate module. After that, GVCF files for each sample were generated using GATK v4.3 with the HaplotypeCaller module ^104^, and variant sites were obtained using GenotypeGVCFs, with SNP/Indel sites obtained using SelectVariants. Finally, low-quality SNPs were filtered using VariantFiltration (QD < 10.0 || FS > 60.0 || MQ < 40.0 || SOR > 3.0 || MQRanksum < -12.5 || ReadPosRanksum < -8.0) and these sites underwent annotation through the utilization of SnpEff software ^105^. Considering the issues of insufficient coverage for low depth and amplification of sequencing errors for extremely high depth, sites with less than one-third of the average depth and more than three times the average depth were filtered out. In addition, structural variant analyses were performed. Using the LUMPY program ^106^ and Manta software ^107^ with default parameters, the BAM files aligned to the Nipponbare reference genome were processed. Subsequently, SURVIVOR ^108^ was employed to merge the results and calculate different types of structural variation features. The density plot of genome-wide variant including SNPs and SVs from elite super-hybrid rice and its parental progenitors were visualized by Circos v0.69-8 (Krzywinski et al. 2009) with 1 Mb sliding windows. Finally, to assess the relative abundance of TE in the genome, we filtered the NGS data generated for the five super hybrid rice and its parental samples in this study using the previous parameters, mapping the filtered high-quality reads to the TE annotated sequence database of Nipponbare (http://omap.org/cgi-bin/rite/index.cgi) ^109^. Subsequently, the number of reads mapped for each type of TE is counted. The relative abundance of each TE family and quantitative interpretation is provided using the scale “log10 (UD),” where UD is defined as the mapping read depth of each TE family divided by the total mapping read depth of all TE families.

### Hybridization and introgression analysis among *Oryza* species

To elucidate gene flow dynamics within rice species and its interactions with wild rice species, we conducted hybridization analyses at both gene and SNPs level. At gene level, we employed a rigorous approach to identify putative single copy orthologs from an extensive dataset of 39,984 rooted multi-copy gene trees by using module Ortho_Retriever from PhyloTracer (https://github.com/YiyongZhao/PhyloTracer). To reduce the potential biases introduced by gene loss, we meticulously curated a gene dataset comprising single-copy gene sets with 100% species coverage across 34 genomic datasets. To investigate the signals of ancient hybridization among the MRCAs of the three-rice subspecies, *indica*, *japonica*, and *aus*, our study utilized 1,383 gene duplicates that originated from the MRCA of rice for subsequent HyDe analyses. The module Ortho_Retriever from PhyloTracer was employed to split these multi-copy gene trees into single-copy gene trees and to pinpoint the orthologous genes. Ultimately, we identified a total of 669 orthologous genes that originated from the MRCA of rice, each exhibiting a minimum taxa coverage of 50%. We separately processed these two sets of genes as follows. First, the protein sequences from these orthologous gene sets were aligned by MAFFT v7.508 ^92^ with parameters “--auto”. These protein alignments were then converted into CDS alignments with the guidance from nucleotide alignments by using PAL2NAL ^93^. The resulting CDS alignments were then concatenated into a supermatrix alignment for hybridization analysis by HyDe ^63^, operating under default parameters with a significance threshold of P-value < 0.05.

In the SNP-level analysis, we conducted genetic analysis between Asian rice species and within *indica* rice using two datasets. The first dataset referred to as Dataset 1, comprised genomic NGS data from previous studies, including Asian cultivated rice and wild rice (Table S8). The second dataset, referred to as Dataset 2, consisted of genomic NGS data from this study, including LYP9, Y1, Y2, Y900, and CY1000, along with their parental lines. The steps were as follows: Firstly, the raw data were filtered and quality-controlled using Trimmomatic and high-quality reads were aligned with the Nipponbare genome using the mem module of the BWA software to generate SAM format files. SAMtools v1.16.1 classified the SAM files as BAM files and tagged for repetitive sequences using the Picard software MarkDuplicate module to mark duplicate sequences. After that, GVCF files were generated for each sample using GATK v4.3 and combine all GVCF files using the “CombineGVCFs” parameter to obtain the variant sites using GenotypeGVCFs and SNP sites using SelectVariants. For these SNP loci use VariantFiltration to filter low-quality SNPs and given that anomalous depths can lead to a widening of the data error, we filtered loci below one-third of the mean depth and above three times the mean depth. Finally, Filter loci with missing data > 0.2 and filter linkage disequilibrium (LD) loci using PLINK with the parameter “--indep-pairwise 50 10 0.2” ^110^ to obtain high-quality SNPs. We concatenated the SNPs (Dataset 1: 35502; Dataset 2: 90113) generated from these two datasets into sequence matrices, respectively, and performed hybridization and introgression analyses using HyDe software ^63^. In addition, due to the existence of datasets that do not contain all types of Asian wild rice species, we used different strategies for our analyses. For dataset 1, we performed genetic network analyses between subspecies using TreeMix v1.13 with parameter “-k 500” ^65^. For dataset 2, we performed intra-subspecies genetic network analyses using HyDe.

### Population structure analysis by ADMIXTURE

To analyze the genealogical structure of super rice at the level of genomic mutations, we performed population structure analysis using ADMIXTURE ^111^ for 90,113 high-quality SNP loci identified in the genomic NGS data of the five samples of super rice and its parents in the present study, with a K-value ranging from 2 to 15. The results were presented graphically using the ggplot2 software package for R. The results are shown below. Afterward, a graphical presentation was performed using the ggplot2 package in R.

### Gene expression quantification and clustering for RNA-seq data

RNA-seq data collected for the three super-hybrid rice varieties (LYP9, Y900, and CY1000) and their hybrid progenitors, tissues including roots, stems, leaves, and panicle fluff, are detailed in Table S11. High-quality reads were obtained by filtering and quality control of raw data using Trimmomatic (v0.39, options: LEADING:10 TRAILING:10 SLIDINGWINDOW:4:20 MINLEN:36) ^101^. The clean reads were then mapped to the *O. sativa* Nipponbare reference genome using HISAT2 (v2.2.1) ^112^, followed by quantification of gene expression using StringTie ^113^. To preserve more information regarding differentially expressed genes, for each gene in the hybrid combinations, comparisons were made between offspring and both parental lines. If a gene exhibited significant differential expression in any comparison, it was retained and the final difference dataset is obtained. The dataset was clustered using the R package ComplexHeatmap ^114^. To facilitate the discovery of gene expression pattern between progeny and parents, all genes were organized into data matrices according to parental combinations and clustering was conducted using the R package Mfuzz ^70^.

### Classification for gene expression patten between progeny and progenitors

Previous studies on LYP9 suggest that non-additive genes may be the dominant factor in dominance formation. To further validate the measurement point of view, we additionally included the transcriptome data of two super rice combinations, Y900 and CY1000, to classify the genes statistically with consistent criteria ^40^. The details are as follows: If the A gene was significantly higher than the higher parent gene, the gene was classified as positive overdominance (POD). If the A gene was significantly lower than the lower parent gene, the gene was classified as negative overdominance (NOD). If A the gene was significantly higher than the middle-parent (The middle-parent expression level is calculated as (P1 + P2)/2, where P1 and P2 represent the expression levels of each gene in both parents.) but showed no significance from the higher parent, the gene was classified as positive dominance (PD). If the A gene was significantly lower than the middle-parent but showed no significance from the lower parent, the gene was classified as negative dominance (ND). If the A gene was significantly higher than the middle-parent and significantly lower than the higher parent, the gene was classified as positive partial dominance (PPD). If the A gene was significantly lower than that of the middle-parent and significantly higher than that of the lower parent, the gene was classified as negative partial dominance (NPD). If the A gene was not significantly different from the middle-parent, significantly lower than the higher parent, and significantly higher than the lower parent, the gene was classified as additive expression gene (A).

### eQTL analysis

In our study, we conducted a linear association analysis to investigate expression quantitative trait loci (eQTLs) across three elite super-hybrid rice varieties (LYP9, Y900, and CY1000) and their hybrid parents, integrating comprehensive gene expression data from their transcriptomes with genomic variant information. Using fastQTL software ^115^, we identified significant eQTL sites by applying a significance threshold of P ≤ 1E^-5^ for Y900 and CY1000, and a relaxed threshold of P ≤ 1E^-4^ for LYP9 (exhibited a lower number of associated sites) to accommodate the identification of high-value signal sites. This analysis revealed 25 genes associated with significant eQTL sites, implicated in a range of biological functions from stress responses to the determination of key plant morphological traits. To elucidate the relationships between phenotypes, genes, and eQTL sites, we generated detailed association diagrams using the ggplot2 package in R, providing a visual representation of the genetic architecture underlying trait variation in these rice varieties.

### KEGG and GO enrichment analysis

In the MRCA of *Oryza sativa*, a comprehensive reconciliation of gene family phylogenies with the overarching species phylogeny revealed a total of 1,383 gene duplication events. We find 158 genes of the *Oryza sativa* Nipponbare genome derived from these duplication events. In addition, we obtained a group of genes significantly associated with the mutant loci in the eQTL analysis. For these above two gene sets, Kyoto Encyclopedia of Genes and Genomes (KEGG) and Gene Ontology (GO) enrichment analysis was performed using the OmicShare tool (https://www.omicshare.com/tools/home/report/koenrich.html).

### Identification of de novo SNPs in elite super-hybrid rices

The methodology for identifying shared genetic sites between progeny and parental genotypes involved treating each hybrid and their progenitors as a discrete unit. Within this framework, a genetic site in the progeny was classified as homozygous for a mutation if it matched the genotype of either parent, indicating inheritance from that parent. Sites consistent with both parents’ genotypes were categorized as shared, whereas sites where the mutant genotype of progeny diverged from both parents were designated as de novo. These sites underwent annotation through the utilization of SnpEff software ^105^ to delineate genes harboring de novo SNPs. Concurrently, gene expression profiles were ascertained by integrating transcriptome data, facilitating a comprehensive understanding of the genetic and functional landscape of these elite super-hybrid rice varieties.

## Statements & Declarations

## Author contributions

Yiyong Zhao and Quanzhi Zhao conceived and designed the experiments. Quanzhi Zhao prepared rice samplings and subsequent sequencing. Daliang Liu performed most of the bioinformatic analyses under the supervision and assistance from Yiyong Zhao. Hao Yin contributed to the curation and preliminary cleaning of the Poaceae genomic data. Tao Li supported the phylogenomic analyses. Liang Wang carried out experiments to test the genetic variant findings. Taikui Zhang helped in experimental materials and data sources. The manuscript was composed by Daliang Liu and Yiyong Zhao. Critical revisions were made by Yiyong Zhao, Quanzhi Zhao, Daliang Liu, Houlin Yu, Hao Yin, Tao Li, Liang Wang, Song Lu, Xinhao Sun and Taikui Zhang. All authors thoroughly reviewed and endorsed the final version of the manuscript.

## Data availability

All data that support the findings of this study are available from the corresponding author upon a reasonable request.

## Ethical approval

The authors declare that the work is original research that has not been published previously and is not under consideration for publication elsewhere.

## Consent to participate

Informed consent was obtained from all individual participants included in the study.

## Acknowledgments

The authors are particularly grateful to the anonymous reviewers for their valuable comments on the manuscript. We are grateful to the State Key Laboratory of Big Data, Guizhou University, for providing supercomputing services for the data analysis. We thank Dr. Jian He from Beijing Forestry University for his assistance in visualization methodology of Hyde.

## Cover image candidate

Here, is an AI-aided cover image, concentrating on the origins and heterosis of the cultivated rice. It artistically depicts the hybridization process of an elite rice hybrid, grounded in a DNA double helix, and set against a backdrop of reticulate evolution, showcasing gene flow and the generation of de novo SNPs.

## Supplementary figure legends

**Figure S1.** A summary of the relative domestication time between subspecies of cultivated rice and the distribution characteristics of Ks density of all homologous gene pairs within the cultivated rice species.

(a) Boxplots showing the Ks values of all syntenic gene pairs between the wild rice and the representative species of two subspecies of cultivated rice, respectively, which are boxplot representations of the corresponding species in Fig. 1d. (b) shows the trend in Ks density for all paralogous gene pairs in the respective genomes of *Oryza sativa* rice varieties, and the right-hand legend shows population information. (c) A molecular clock analysis based on concatenated sequences of the 54 putative domestication genes identified from low nucleotide diversity regions, illustrating the relative timing of domestication across subspecies of cultivated rice. The blue bars represent the 95% highest posterior density (HPD) for the estimated divergence times. The pentagrams represent the MRCA of each rice subgroup, with divergence times denoted to the right. The number of gene duplication events is indicated at the upper left of each node, with the red star marking the MRCA of *Oryza sativa*.

**Figure S2.** Genomic visualization of gene duplications originating from the MRCA of *Oryza sativa* by Circos and dot plots.

This figure delineates the gene duplication dynamics within the MRCA of *Oryza sativa* by using *japonica* (Nipponbare, left panels) and *indica* (R9311, right panels) subpopulations as representatives. Subfigures (a) and (b) display the collinearity among duplicated genes across the rice genome. Subfigures (c) and (d) present a genome-wide mapping of synonymous substitution rates (Ks) for the Nipponbare and R9311 genome, indicating the absence of recent whole-genome duplication (WGD) events in the ancestors of genus *Oryza*. Subfigures (e) and illustrate the chromosomal distribution of duplicated gene pairs within subspecies of the Asian cultivated rice, highlighting the positional distribution of genomic regions of gene duplication events, 402 and 81 duplicated gene pairs were involved in Nipponbare and R9311, respectively.

**Figure S3.** Comparative analysis of Ka, Ks, and Ka/Ks ratios for 11 key domestication genes in *japonica* and *indica* subspecies.

This figure summarizes the non-synonymous (Ka), synonymous (Ks), and the ratio of non-synonymous to synonymous substitutions (Ka/Ks) for eleven crucial genes associated with the domestication of *japonica* and *indica* rice subspecies. The histogram subpanels represent the variation in these values across the *japonica* and *indica* comparisons with different wild and cultivated rice varieties. Each bar color correlates with a specific comparison as indicated in the legend, facilitating the assessment of divergence and evolutionary pressures exerted on these domestication-related genes.

**Figure S4.** Hybridization and introgression signal for the MRCAs of *O. sativa* and three rice subspecies, respectively.

The heatmap displays hybridization signals as detected by HyDe for each MRCAs of *Oryza sativa*, *indica*, *japonica*, and *aus*, respectively, annotated with red-highlighted branches on the cladogram to denote the detected clade. The analysis was designed to incorporate two distinct gene sets of single-copy gene. The first gene set, consisting of 669 genes, covering 50% of Asian rice taxa; These genes have been identified as originating from duplication events in the MRCA of *O. sativa*, and the purpose for this analysis is to analyze the ancient hybridization among different MRCAs of *indica*, *japonica* and *aus* (a, c, e and g). The other gene set comprised 1,300 orthologous genes covering 100% rice taxa (b, d, f and h). These two gene sets collectively facilitate hybridization analyses of the ancestral hybridization among different rice subspecies, as well as providing insights into the more recent genetic hybridization events. See methods for detailed gene selection procedures. Subfigures (a) and (b) focus on the progenitors of Asian cultivated rice, whereas (c) and (d) concentrate on the *indica* lineage, (e) and (f) on the *japonica* lineage, and (g) and (h) on the *aus* lineage. Each small colored square on the heatmap corresponds to a hybridization signal identified by HyDe, with the color intensity representing the probability of inheritance for the associated taxa on the y-axis. Contiguous squares of identical shading that compose a larger block within the heatmap, along with their corresponding taxa on the axes forming a monophyletic clade, which is indicated with a node number on a green background.

**Figure S5.** Genomic SVs landscape of five super-hybrid rice and their hybrid progenitors.

The subfigures (a-e) depict the landscape of structural variations for deletion, duplication, inversion, insertion, and translocation, respectively. The Circos plots represent the SV density within the genomes of five super-hybrid rice varieties and their progenitors, mapped using a 1000 kb sliding window approach. The outermost ring (a) represents 12 rice chromosomes, marked in Mb units. The second ring (b) illustrates gene density. Rings (c-n) represent SV densities for the rice verities: LYP9, PA64S, Y58S, R9311, Y1, Y2, YH2, R900, Y900, GX24S, CY1000, and *O. rufipogon*, respectively.

**Figure S6.** Distribution of genomic variations (SNPs and InDels) in different genomic regions of five super-hybrid rice varieties and their parental progenitors.

The figure illustrates the distribution of SNPs and InDels across various genomic regions; specifically, 2 kilobases (kb) upstream and downstream of genes, exons, introns, and non-coding regions in five super-hybrid rice varieties alongside their respective hybrid parents. Panel (a) quantifies the ratio of SNPs, while panel (b) details the ratio of InDels, both of which are critical indicators of genetic diversity and evolution within these varieties. The data are represented as stacked bar graphs, with color coding to differentiate between the genomic regions.

**Figure S7.** Distribution of SVs in genomic regions of five super-hybrid rice varieties and their parental progenitors.

Panel a and b of Figure S7 depict the distribution of SVs, including deletions (DEL), duplications (DUP), insertions (INS), inversions (INV), and translocations (TRA), across different genomic regions of five super-hybrid rice varieties and their respective parental progenitors. These regions encompass two kilobases (kb) upstream and downstream of genes, as well as exons, introns, and intergenic regions. The data are represented in stacked bar charts for each sample, with colors indicating the type of SVs.

**Figure S8.** Comparative heatmap of summary of TE abundance in the genomes of five super-hybrid rice varieties and their progenitors, Nipponbare, and Asian wild rice.

This heatmap provides a comparative summary of the relative abundance of various TE families within the genomes of eleven *indica* rice varieties, the *japonica* variety Nipponbare, and Asian wild rice. The rows of the heatmap correspond to different TE families, while the columns represent individual rice varieties and species. The color gradient reflects the relative abundance of each TE family, and quantitative interpretation is provided using the scale “log10(UD),” where UD is defined as the mapping read depth of each TE family divided by the total mapping read depth of all TE families.

**Figure S9.** Summary of the hybridization signal (γ) from HyDe analysis among the five super-hybrid rice varieties and their parental progenitors.

Heatmap displays the hybridization signals detected by HyDe among five sets of super-hybrid rice and their parental progenitors, as well as the Nipponbare, with *O. rufipogon* as an outgroup. Each colored square on the heatmap corresponds to a hybridization signal identified by HyDe, with the intensity of color representing the genetic probability of the associated taxa on the Y-axis.

**Figure S10.** A Venn diagram summarizing overlapping eQTL loci among three super-rice varieties (LYP9, Y900, and CY1000) and their parental progenitors.

**Figure S11.** The bubble plot summarizing the biological processes of eQTL genes for three super-hybrid rice varieties and their parental progenitors based on Gene Ontology (GO). The figure summarizes the annotation of important biological processes (BP) terms (P-value < 0.05) for eQTL genes in the LYP9, Y900, and CY1000 super-hybrid rice varieties and their parental progenitors based on the Gene Ontology (GO) database.

**Figure S12.** The bubble plot summarizes the KEGG pathway annotations (P-value < 0.05) for eQTL genes within the LYP9, Y900, and CY1000 super-hybrid rice varieties and their parental progenitors.

## Supplementary tables legends

**Table S1.** The information of the genomic datasets employed for phylogeny reconstruction, encompassing 34 genomes including 33 genomes from Oryzeae and *Brachypodium distachyon* as outgroup.

**Table S2.** Summary of Gene Ontology (GO) terms (Q-value < 0.05) for 158 genes of the *Oryza sativa* Nipponbare genome derived from duplication events in the MRCA of *Oryza sativa* ancestors aims to explain the impact of duplication events on the *Oryza sativa* Nipponbare.

**Table S3.** Summary of Kyoto Encyclopedia of Genes and Genomes (KEGG) pathways (Q-value < 0.05) for 158 genes of the *Oryza sativa* Nipponbare genome derived from duplication events in the MRCA of *Oryza sativa* aims to explain the impact of duplication events on the species within the *Oryza sativa* Nipponbare.

**Table S4.** Summary of divergence time estimation of 54 putative domesticated genes. These genes were identified in 15 regions of low genomic nucleotide diversity. The MRCA of *O. sativa*, *japonica*, *indica*, and *aus* in the table correspond to nodes of species tree in Fig. 1, the time unit: million years ago (Mya).

**Table S5.** Summary of Ks (synonymous substitution rate) values for 54 orthologous gene pairs of putatively domesticated genes, identified across 15 genomic regions exhibiting low nucleotide diversity. To approximate the onset of domestication process for various cultivated rice ancestors, we employed multiple representative genomes from different *Oryza* subgroups. This was done to calculate the Ks values for those 54 orthologous domesticated genes, thereby providing an estimation of the origin of domestication process. The domestication origin of MRCA of *O. sativa* was inferred from a comparison between *O. rufipogon* w1943 and *O. rufipogon* w1654. The domestication origin of MRCA of *japonica* inferred from a comparison between *O. rufipogon* w1943 and *O. sativa* Nipponbare. The domestication origin of the MRCA of *indica* was inferred from a comparison between *O. rufipogon* w1654 and *O. sativa* R9311. Lastly, the domestication origin of the MRCA of *aus* was inferred from a comparison between *O. nivara* and *O. sativa aus* n22.

**Table S6.** Proportional distribution of gene duplication types in ancestral nodes of cultivated rice, detailing the prevalence of tandem duplication, genomic collinearity, and other forms of gene duplication. The analysis is based on the *O. sativa* Nipponbare and *O. sativa* R9311 genomes, which serve as representative species for the *japonica* and *indica* subgroups, respectively.

**Table S7.** Chi-square test was performed on the TE annotated within the *japonica* representative Nipponbare and the *indica* representative R9311 vs. duplicated genes. TE genes refer to genes located within the 2 kb upstream and downstream of TE regions.

**Table S8.** The details of the whole genome sequencing datasets downloaded from the European Nucleotide Archive (ENA) for TreeMix analysis.

**Table S9.** Detailed quality assessment of newly sequenced whole genome sequencing data preprocessing results for five super-hybrid rice varieties and their parental progenitors in this study.

**Table S10.** Quantification of SNPs, InDels, and SVs detected in newly sequenced whole genome sequencing data for five super-hybrid rice varieties with their parental progenitors and *Oryza rufipogon* in this study.

**Table S11.** Quantification on the classification of genomic variants and their distribution in different rice varieties from newly sequenced whole genome sequencing data in this study.

**Table S12.** Summary of the heritability between five super rice hybrids and their parents, as well as Nipponbare. Based on hybrid analysis using 90,113 SNP loci, where P1 and P2 represent the maternal and paternal parents, respectively, and Gamma indicates the genetic contribution of P1 to the hybrid.

**Table S13.** The information of RNA sequencing data for the three super-hybrid rice varieties (LYP9, Y900, and CY100) with their parental progenitors.

**Table S14.** Summary of annotation information for genes related to yield traits from the China Rice Data Center database (https://www.ricedata.cn/).

**Table S15.** Summary of gene expression profiles across three super-hybrid rice varieties and their progenitors. This table presents a comprehensive overview of gene expression data collected from three super-hybrid rice varieties: LYP9, Y900, and CY1000. It also includes data from their progenitors: GX24S, PA64s, R900, R9311, and Y58S. The data encompasses gene expression levels in different plant tissues, specifically leaves, stems, and panicles, offering insights into the gene expression dynamics across various stages of plant growth and development in both the hybrid varieties and their ancestral lines.

**Table S16.** Differential gene expression clustering in super-hybrid rice varieties and their progenitors. This table delineates the results of gene expression clustering using the MFUZZ algorithm for three super-hybrid rice varieties, namely LYP9, Y900, and CY1000, along with their progenitor strains. Displayed within the table are clusters of differentially expressed genes. Each gene’s expression data is associated with a specific tissue sample, using a naming convention that includes the gene identifier, variety code, and tissue type, separated by underscores.

**Table S17.** Summary of the gene expression patterns in various tissue samples. ’M_’ indicates genes expressed from the paternal side, ’F_’ designates maternal gene expression, and ’F1_*’ highlights the gene expression in the hybrid progeny. The patterns of gene expression are succinctly coded as follows: POD for positive overdominance, NOD for negative overdominance, PD for positive dominance, ND for negative dominance, PPD for positive partial dominance, NPD for negative partial dominance, and A for additive expression. For comprehensive definitions of these terms, readers are directed to consult the Methods section of the document.

**Table S18.** Summary of eQTL genes identified for three super-hybrid rice varieties and their progenitors. This table provides a compilation of eQTL genes that have been identified in three super-hybrid rice varieties, namely Y900 and CY1000, employing a significance threshold of P-value < 1E^-5^, and for LYP9 with a relaxed threshold of P-value < 1E^-4^.

**Table S19.** Trait ontology annotations for marker genes with significant eQTL signals. This table provides detailed trait ontology annotations for marker genes that exhibit strong eQTL signals, aiding in the elucidation of genetic influences on specific traits. The associated trait information for each gene was sourced from The Rice Annotation Project (RAP, http://rice.uga.edu/).

**Table S20.** Gene Ontology (GO) enrichment analysis for eQTL genes specifically expressed in LYP9.

**Table S21.** GO enrichment analysis for eQTL genes specifically expressed in Y900.

**Table S22.** GO enrichment analysis for eQTL genes specifically expressed in CK1000.

**Table S23.** KEGG pathway enrichment analysis of eQTL genes in three super-hybrid rice varieties (LYP9, Y900, and CY1000).

**Table S24.** Summary of yield-related genes linked to de novo SNP loci in four super-hybrid rice varieties. This table compiles a list of yield-related genes that are associated with de novo single nucleotide polymorphism (SNP) loci identified in four super rice varieties: Y1, Y2, Y900, and CY1000. Notably, these specific SNP loci have not been detected in the LYP9 variety.

**Table S25.** Characterization of trait-associated genes with de novo SNP loci and expression profiles. This table outlines the traits of genes linked with de novo SNP (single nucleotide polymorphism) loci as indicated in Figure 7a, detailing the gene IDs, gene names, chromosomal positions, gene start and end points, and trait descriptions. Genes that are actively expressed are highlighted in red. Additionally, the table includes information on where these genes are expressed across three super-hybrid rice varieties LYP9, Y900, and CY1000.

## References

1. Vaughan, D. A., Morishima, H. & Kadowaki, K. Diversity in the *Oryza* genus. Curr. Opin. Plant Biol. 6, 139–146 (2003).

2. Khush, G. S. Origin, dispersal, cultivation and variation of rice. Plant Mol.Biol. 35, 25–34 (1997).

3. Balakrishnan, D. et al. Detecting CSSLs and yield QTLs with additive, epistatic and QTL×environment interaction effects from Oryza sativa × O. nivara IRGC81832 cross. Sci Rep. 10, 7766 (2020).

4. Dong, L. et al. Identification and Fine Mapping of Pi69(t), a New Gene Conferring Broad-Spectrum Resistance Against Magnaporthe oryzae From Oryza glaberrima Steud. Front. Plant Sci. 11, 1190 (2020).

5. Swamy, B. P. & Sarla, N. Yield-enhancing quantitative trait loci (QTLs) from wild species. Biotechnol. Adv. 26, 106–120 (2008).

6. Garris, A. J., Tai, T. H., Coburn, J., Kresovich, S. & McCouch, S. Genetic structure and diversity in *Oryza sativa* L. Genetics. 169, 1631–1638 (2005).

7. Hour, A. L. et al. Genetic Diversity of Landraces and Improved Varieties of Rice (*Oryza sativa* L.) in Taiwan. Rice. 13, 82 (2020).

8. Gaikwad, K. B. et al. Deployment of wild relatives for genetic improvement in rice (*Oryza sativa*). Plant Breed. 140, 23–52 (2021).

9. Gross, B. L. & Zhao, Z. Archaeological and genetic insights into the origins of domesticated rice. Proc. Natl. Acad. Sci. U. S. A. 111, 6190–6197 (2014).

10. Civan, P., Craig, H., Cox, C. J. & Brown, T. A. Three geographically separate domestications of Asian rice. Nat. Plants. 1, 15164 (2015).

11. Bates, J., Petrie, C. A. & Singh, R. N. Approaching rice domestication in South Asia: New evidence from Indus settlements in northern *India*. J. Archaeol. Sci. 78, 193–201 (2017).

12. Choi, J. Y. et al. The Rice Paradox: Multiple Origins but Single Domestication in Asian Rice. Mol. Biol. Evol. 34, 969–979 (2017).

13. Ghazy, M. I., Salem, K. & Sallam, A. Utilization of genetic diversity and marker-trait to improve drought tolerance in rice (*Oryza sativa* L.). Mol. Biol. Rep. 48, 157–170 (2021).

14. Sasaki, T. The map-based sequence of the rice genome. Nature. 436, 793–800 (2005).

15. Yang, G. et al. Genetic Diversity of Shanlan Upland Rice (*Oryza sativa* L.) and Association Analysis of SSR Markers Linked to Agronomic Traits. Biomed Res. Int. 2021, 7588652 (2021).

16. Zhu, B. F. et al. Genetic control of a transition from black to straw-white seed hull in rice domestication. Plant Physiol. 155, 1301–1311 (2011).

17. Tan, L. et al. Control of a key transition from prostrate to erect growth in rice domestication. Nature Genet. 40, 1360–1364 (2008).

18. Molina, J. et al. Molecular evidence for a single evolutionary origin of domesticated rice. Proc. Natl. Acad. Sci. U. S. A. 108, 8351–8356 (2011).

19. Huang, X. et al. A map of rice genome variation reveals the origin of cultivated rice. Nature. 490, 497–501 (2012).

20. Yang, C. C. et al. Independent domestication of Asian rice followed by gene flow from japonica to indica. Mol. Biol. Evol. 29, 1471–1479 (2012).

21. Wambugu, P. W., Brozynska, M., Furtado, A., Waters, D. L. & Henry, R. J. Relationships of wild and domesticated rices (*Oryza* AA genome species) based upon whole chloroplast genome sequences. Sci Rep. 5, 13957 (2015).

22. Jing, C. Y. et al. Multiple domestications of Asian rice. Nat. Plants. 9, 1221–1235 (2023).

23. Hochholdinger, F. & Baldauf, J. A. Heterosis in plants. Curr. Biol. 28, R1089–R1092 (2018).

24. Liu, W., Zhang, Y., He, H., He, G. & Deng, X. W. From hybrid genomes to heterotic trait output: Challenges and opportunities. Curr. Opin. Plant Biol. 66, 102193 (2022).

25. Lu, C. & Zhou, J. Breeding and utilization of two-line interspecific hybrid rice Liangyou Peijiu. Hybrid Rice. 15, 4–5 (2000).

26. Jiang, Z. Study on Individual Plant Type Character of Liangyoupeijiu Rice. Acta Agronomica Sinica, (2003).

27. Deng, Q. Breeding of the PTGMS Line Y58S with Wide Adaptability in Rice. Hybrid Rice, (2005).

28. Liao, F. The phase Ⅱ target of the chinese single cropping super rice realized one year ahead of schedule (In China). Hybrid Rice. 06, 102 (2004).

29. Wu, J., Deng, Q., Yuan, D. & Qi, S. Progress of super hybrid rice research in China. Chinese Science Bulletin. 61, 3787–3796 (2016).

30. Chang, S. et al. Studies on Matching Technology of “Four Elite Factors” for Super Hybrid Rice with Yield 15.0 t/hm2. China Rice. 21, 1–6 (2015).

31. Xin, Y. New Research Progress Achieved for Super Hybrid Rice Chaoyou 1000 (in China). Hybrid Rice. 35(**2**), 102 (2020).

32. Bruce, A. B. The Mendelian Theory of Heredity and the Augmentation of Vigor. Science. 32, 627–628 (1910).

33. East, E. M. Heterosis. Genetics. 21, 375–397 (1936).

34. Minvielle, F. Dominance is not necessary for heterosis: a two-locus model. Genet. Res. 49, 245–247 (1987).

35. Krieger, U., Lippman, Z. B. & Zamir, D. The flowering gene *SINGLE FLOWER TRUSS* drives heterosis for yield in tomato. Nature Genet. 42, 459–463 (2010).

36. Swanson-Wagner, R. A. et al. All possible modes of gene action are observed in a global comparison of gene expression in a maize F1 hybrid and its inbred parents. Proc. Natl. Acad. Sci. U. S. A. 103, 6805–6810 (2006).

37. Li, X., Li, X., Fridman, E., Tesso, T. T. & Yu, J. Dissecting repulsion linkage in the dwarfing gene *Dw3* region for sorghum plant height provides insights into heterosis. Proc. Natl. Acad. Sci. U. S. A. 112, 11823–11828 (2015).

38. Huang, X. et al. Genomic architecture of heterosis for yield traits in rice. Nature. 537, 629–633 (2016).

39. Shao, L. et al. Patterns of genome-wide allele-specific expression in hybrid rice and the implications on the genetic basis of heterosis. Proc. Natl. Acad. Sci. U. S. A. 116, 5653–5658 (2019).

40. Xie, J. et al. Large-scale genomic and transcriptomic profiles of rice hybrids reveal a core mechanism underlying heterosis. Genome Biol. 23, 264 (2022).

41. Wang, X. et al. Genome-wide analysis of transcriptional variability in a large maize-teosinte population. Mol. Plant. 11, 443–459 (2018).

42. Wang, W. et al. Genomic variation in 3,010 diverse accessions of Asian cultivated rice. Nature. 557, 43–49 (2018).

43. Yoshida, H. et al. Genome-wide association study identifies a gene responsible for temperature-dependent rice germination. Nat. Commun. 13, 5665 (2022).

44. Adhikari, K. et al. A genome-wide association scan in admixed Latin Americans identifies loci influencing facial and scalp hair features. Nat. Commun. 7, 10815 (2016).

45. Vosa, U. et al. Large-scale cis- and trans-eQTL analyses identify thousands of genetic loci and polygenic scores that regulate blood gene expression. Nature Genet. 53, 1300–1310 (2021).

46. Pang, J. et al. Kernel size-related genes revealed by an integrated eQTL analysis during early maize kernel development. Plant J. 98, 19–32 (2019).

47. Chen, D., Zhang, T., Chen, Y., Ma, H. & Qi, J. Tree2GD: a phylogenomic method to detect large-scale gene duplication events. Bioinformatics. 38, 5317–5321 (2022).

48. Kombrink, E. & Somssich, I. E. Defense Responses of Plants to Pathogens. In: Callow, J. A., Andrews, J. H. & Tommerup, I. C., editors. Advances in Botanical Research: Academic Press; 1995. pp. 1–34.

49. Zhang, L., Du L & Poovaiah, B. W. Calcium signaling and biotic defense responses in plants. Plant Signal. Behav. 9, e973818 (2014).

50. Wan, W. L., Frohlich, K., Pruitt, R. N., Nurnberger, T. & Zhang, L. Plant cell surface immune receptor complex signaling. Curr. Opin. Plant Biol. 50, 18–28 (2019).

51. Lea, C. S., Bradbury, S. G. & Constabel, C. P. Anti-Herbivore Activity of Oregonin, a Diarylheptanoid Found in Leaves and Bark of Red Alder (*Alnus rubra*). J. Chem. Ecol. 47, 215–226 (2021).

52. Doebley, J. F., Gaut, B. S. & Smith, B. D. The molecular genetics of crop domestication. Cell. 127, 1309–1321 (2006).

53. Konishi, S. et al. An SNP caused loss of seed shattering during rice domestication. Science. 312, 1392–1396 (2006).

54. Kuroha, T. et al. Ethylene-gibberellin signaling underlies adaptation of rice to periodic flooding. Science. 361, 181–186 (2018).

55. Gu, B. et al. *An-2* Encodes a Cytokinin Synthesis Enzyme that Regulates Awn Length and Grain Production in Rice. Mol. Plant. 8, 1635–1650 (2015).

56. Duan, P. et al. Natural Variation in the Promoter of *GSE5* Contributes to Grain Size Diversity in Rice. Mol. Plant. 10, 685–694 (2017).

57. Wang, Y. & Li, J. Rice, rising. Nature Genet. 40, 1273–1275 (2008).

58. Sun, X. et al. The C-S-A gene system regulates hull pigmentation and reveals evolution of anthocyanin biosynthesis pathway in rice. J. Exp. Bot. 69, 1485–1498 (2018).

59. Zhao, B. et al. *Sdr4* dominates pre-harvest sprouting and facilitates adaptation to local climatic condition in Asian cultivated rice. J. Integr. Plant Biol. 64, 1246–1263 (2022).

60. Yan, H. et al. Multiple tissue-specific expression of rice seed-shattering gene *SH4* regulated by its promoter pSH4. Rice. 8, 12 (2015).

61. Zhou, Y. et al. Genetic control of seed shattering in rice by the APETALA2 transcription factor *SHATTERING ABORTION1*. Plant Cell. 24, 1034–1048 (2012).

62. Sato, H., Suzuki, Y., Sakai, M. & Imbe, T. Molecular Characterization of Wx-mq, a Novel Mutant Gene for Low-amylose Content in Endosperm of Rice (Oryza sativa L.). Breed. Sci. 52, 131–135 (2002).

63. Blischak, P. D., Chifman, J., Wolfe, A. D. & Kubatko, L. S. HyDe: A Python Package for Genome-Scale Hybridization Detection. Syst. Biol. 67, 821–829 (2018).

64. Stein, J. C. et al. Genomes of 13 domesticated and wild rice relatives highlight genetic conservation, turnover and innovation across the genus *Oryza*. Nature Genet. 50, 285–296 (2018).

65. Pickrell, J. K. & Pritchard, J. K. Inference of population splits and mixtures from genome-wide allele frequency data. Plos Genet. 8, e1002967 (2012).

66. Evanno, G., Regnaut, S. & Goudet, J. Detecting the number of clusters of individuals using the software STRUCTURE: a simulation study. Mol. Ecol. 14, 2611–2620 (2005).

67. Mei, H. et al. Experimental validation of inter-subspecific genetic diversity in rice represented by the differences between the DNA sequences of ’Nipponbare’ and ’93-11’. Chinese Science Bulletin. 52, 1327–1337 (2007).

68. Sun, Z. et al. Dissecting the genetic basis of heterosis in elite super-hybrid rice. Plant Physiol. 192, 307–325 (2023).

69. Patterson, N. et al. Ancient admixture in human history. Genetics. 192, 1065–1093 (2012).

70. Kumar, L. & E, F. M. Mfuzz: a software package for soft clustering of microarray data. Bioinformation. 2, 5–7 (2007).

71. Baker, R. L. et al. Integrating transcriptomic network reconstruction and eQTL analyses reveals mechanistic connections between genomic architecture and Brassica rapa development. Plos Genet. 15, e1008367 (2019).

72. Xu, C. et al. Integration of eQTL and GWAS analysis uncovers a genetic regulation of natural ionomic variation in Arabidopsis. Plant Cell Reports. 42, 1473–1485 (2023).

73. Sharma, S., Kaur, C., Singla-Pareek, S. L. & Sopory, S. K. OsSRO1a Interacts with RNA Binding Domain-Containing Protein (OsRBD1) and Functions in Abiotic Stress Tolerance in Yeast. Front. Plant Sci. 7, 62 (2016).

74. Xia-Yu, G. et al. Comparative transcriptomic analysis of the super hybrid rice Chaoyouqianhao under salt stress. Bmc Plant Biol. 22, 233 (2022).

75. Kan, Y. et al. TT2 controls rice thermotolerance through SCT1-dependent alteration of wax biosynthesis. Nat. Plants. 8, 53–67 (2022).

76. Huang, L., et al. Stress-Responsive Expression, Subcellular Localization and Protein-Protein Interactions of the Rice Metacaspase Family. Int. J. Mol. Sci. 16, 16216–16241 (2015).

77. Su, C. F. et al. A novel *MYBS3*-dependent pathway confers cold tolerance in rice. Plant Physiol. 153, 145–158 (2010).

78. Ito, Y. et al. Fatty acid elongase is required for shoot development in rice. Plant J. 66, 680–688 (2011).

79. Wang, H. et al. Rice *WRKY4* acts as a transcriptional activator mediating defense responses toward Rhizoctonia solani, the causing agent of rice sheath blight. Plant Mol.Biol. 89, 157–171 (2015).

80. Huang, P. et al. Phylogeography of Asian wild rice, *Oryza rufipogon*: a genome-wide view. Mol. Ecol. 21, 4593–4604 (2012).

81. Wang, H., Vieira, F. G., Crawford, J. E., Chu, C. & Nielsen, R. Asian wild rice is a hybrid swarm with extensive gene flow and feralization from domesticated rice. Genome Res. 27, 1029–1038 (2017).

82. Vaughan, D. A., Lu, B. & Tomooka, N. The evolving story of rice evolution. Plant Sci. 174, 394–408 (2008).

83. Zhang, C. & Mirarab, S. ASTRAL-Pro 2: ultrafast species tree reconstruction from multi-copy gene family trees. Bioinformatics. 38, 4949–4950 (2022).

84. Cai, Z. et al. Parallel Speciation of Wild Rice Associated with Habitat Shifts. Mol. Biol. Evol. 36, 875–889 (2019).

85. Cheng, L., Kim, K. W. & Park, Y. J. Evidence for selection events during domestication by extensive mitochondrial genome analysis between japonica and indica in cultivated rice. Sci Rep. 9, 10846 (2019).

86. Liu, R., Zheng, X. M., Zhou, L., Zhou, H. F. & Ge, S. Population genetic structure of Oryza rufipogon and Oryza nivara: implications for the origin of O. nivara. Mol. Ecol. 24, 5211–5228 (2015).

87. Wu, J., Guo, X., Peng, Y. & Zhang, Y. High-yielding Cultivation Techniques of Super Hybrid Rice Xiangliangyou 900 with Yield 16.35 t/hm2 (in Chinese). China Rice. 26, 61 (2020).

88. Chatterjee, N. & Walker, G. C. Mechanisms of DNA damage, repair, and mutagenesis. Environ. Mol. Mutagen. 58, 235–263 (2017).

89. Scharer, O. D. Nucleotide excision repair in eukaryotes. Cold Spring Harbor Perspect. Biol. 5, a12609 (2013).

90. Li, X. & Heyer, W. D. Homologous recombination in DNA repair and DNA damage tolerance. Cell Res. 18, 99–113 (2008).

91. Emms, D. M. & Kelly, S. OrthoFinder: phylogenetic orthology inference for comparative genomics. Genome Biol. 20, 238 (2019).

92. Katoh, K., Misawa, K., Kuma, K. I. & Miyata, T. MAFFT: a novel method for rapid multiple sequence alignment based on fast Fourier transform. Nucleic Acids Res. 30, 3059–3066 (2002).

93. Suyama, M., Torrents, D. & Bork, P. PAL2NAL: robust conversion of protein sequence alignments into the corresponding codon alignments. Nucleic Acids Res. 34, W609–W612 (2006).

94. Nguyen, L., Schmidt, H. A., von Haeseler, A. & Minh, B. Q. IQ-TREE: A Fast and Effective Stochastic Algorithm for Estimating Maximum-Likelihood Phylogenies. Mol. Biol. Evol. 32, 268–274 (2015).

95. Dos Reis, M. & Yang, Z. MCMCTree tutorials.; 2017.

96. Buchfink, B., Xie, C. & Huson, D. H. Fast and sensitive protein alignment using DIAMOND. Nat. Methods. 12, 59–60 (2015).

97. Wang, Y. et al. MCScanX: a toolkit for detection and evolutionary analysis of gene synteny and collinearity. Nucleic Acids Res. 40, e49 (2012).

98. Wang, D., Zhang, Y., Zhang, Z., Zhu, J. & Yu, J. KaKs_Calculator 2.0: a toolkit incorporating gamma-series methods and sliding window strategies. Genom. Proteomics Bioinformatics. 8, 77–80 (2010).

99. Wickham, H. ggplot2. Wires Computational Statistics. 3, 180–185 (2011).

100. Storer, J., Hubley, R., Rosen, J., Wheeler, T. J. & Smit, A. F. The Dfam community resource of transposable element families, sequence models, and genome annotations. Mob. Dna. 12, 2 (2021).

101. Bolger, A. M., Lohse, M. & Usadel, B. Trimmomatic: a flexible trimmer for Illumina sequence data. Bioinformatics. 30, 2114–2120 (2014).

102. Jung, Y. & Han, D. BWA-MEME: BWA-MEM emulated with a machine learning approach. Bioinformatics. 38, 2404–2413 (2022).

103. Li, H. et al. The Sequence Alignment/Map format and SAMtools. Bioinformatics. 25, 2078–2079 (2009).

104. McKenna, A. et al. The Genome Analysis Toolkit: a MapReduce framework for analyzing next-generation DNA sequencing data. Genome Res. 20, 1297–1303 (2010).

105. Cingolani, P. et al. A program for annotating and predicting the effects of single nucleotide polymorphisms, SnpEff: SNPs in the genome of Drosophila melanogaster strain w1118; iso-2; iso-3. Fly. 6, 80–92 (2012).

106. Layer, R. M., Chiang, C., Quinlan, A. R. & Hall, I. M. LUMPY: a probabilistic framework for structural variant discovery. Genome Biol. 15, R84 (2014).

107. 107. Chen, X., et al. Manta: rapid detection of structural variants and indels for germline and cancer sequencing applications. Bioinformatics. 32, 1220–1222 (2016).

108. Jeffares, D. C. et al. Transient structural variations have strong effects on quantitative traits and reproductive isolation in fission yeast. Nat. Commun. 8, 14061 (2017).

109. Copetti, D. et al. RiTE database: a resource database for genus-wide rice genomics and evolutionary biology. Bmc Genomics. 16, 538 (2015).

110. Purcell, S. et al. PLINK: a tool set for whole-genome association and population-based linkage analyses. Am. J. Hum. Genet. 81, 559–575 (2007).

111. Alexander, D. H., Novembre, J. & Lange, K. Fast model-based estimation of ancestry in unrelated individuals. Genome Res. 19, 1655–1664 (2009).

112. Kim, D., Paggi, J. M., Park, C., Bennett, C. & Salzberg, S. L. Graph-based genome alignment and genotyping with HISAT2 and HISAT-genotype. Nat. Biotechnol. 37, 907–915 (2019).

113. Pertea, M., Kim, D., Pertea, G. M., Leek, J. T. & Salzberg, S. L. Transcript-level expression analysis of RNA-seq experiments with HISAT, StringTie and Ballgown. Nat. Protoc. 11, 1650–1667 (2016).

114. Gu, Z., Eils, R. & Schlesner, M. Complex heatmaps reveal patterns and correlations in multidimensional genomic data. Bioinformatics. 32, 2847–2849 (2016).

115. Ongen, H., Buil, A., Brown, A. A., Dermitzakis, E. T. & Delaneau, O. Fast and efficient QTL mapper for thousands of molecular phenotypes. Bioinformatics. 32, 1479–1485 (2016).

